# Practical Aspects of Phylogenetic Network Analysis Using PhyloNet

**DOI:** 10.1101/746362

**Authors:** Zhen Cao, Xinhao Liu, Huw A. Ogilvie, Zhi Yan, Luay Nakhleh

## Abstract

Phylogenetic networks extend trees to enable simultaneous modeling of both vertical and horizontal evolutionary processes. PhyloNet is a software package that has been under constant development for over 10 years and includes a wide array of functionalities for inferring and analyzing phylogenetic networks. These functionalities differ in terms of the input data they require, the criteria and models they employ, and the types of information they allow to infer about the networks beyond their topologies. Furthermore, PhyloNet includes functionalities for simulating synthetic data on phylogenetic networks, quantifying the topological differences between phylogenetic networks, and evaluating evolutionary hypotheses given in the form of phylogenetic networks.

In this paper, we use a simulated data set to illustrate the use of several of PhyloNet’s functionalities and make recommendations on how to analyze data sets and interpret the results when using these functionalities. All inference methods that we illustrate are incomplete lineage sorting (ILS) aware; that is, they account for the potential of ILS in the data while inferring the phylogenetic network. While the models do not include gene duplication and loss, we discuss how the methods can be used to analyze data in the presence of polyploidy.

The concept of species is irrelevant for the computational analyses enabled by PhyloNet in that species-individuals mappings are user-defined. Consequently, none of the functionalities in PhyloNet deals with the task of species delimitation. In this sense, the data being analyzed could come from different individuals within a single species, in which case population structure along with potential gene flow is inferred (assuming the data has sufficient signal), or from different individuals sampled from different species, in which case the species phylogeny is being inferred.

## 1 Introduction

Several evolutionary processes acting within and across species can give rise to non-treelike evolutionary histories. These processes include admixture, gene flow, hybridization, hybrid speciation, introgression, and horizontal gene transfer (see Section 7 for details related to PhyloNet and these processes). When any of these processes occurs, the evolutionary history (of the species or sub-populations) is more adequately represented by a network, rather than a tree, structure. In our context, a phylogenetic network takes the shape of a rooted, directed, acyclic graph [21], so that the single root represents the most recent common ancestor of all taxa at the leaves, and every internal node represents either divergence or reticulation.

As reticulation events occur in evolutionary settings—often very closely related species [18], it is important to account simultaneously for *incomplete lineage sorting* (ILS) which also arises under similar conditions. In other words, for accurate inference of reticulate evolutionary histories, gene tree heterogeneity should not be explained solely by reticulation; rather, the potential for ILS to have occurred must be accounted for as well. This is a fundamental assumption in PhyloNet [39], which is a software package that implements a variety of methods for phylogenetic network inference while accounting for both reticulation and ILS.

For parsimonious inference where a network is inferred to minimize a measure of conflict with a set of gene trees, the ‘*minimizing deep coalescence*,’ or MDC, criterion [16, 17, 32] was extended so that it works on species networks [40]. Inference based on this criterion requires a set of gene trees, one per locus (multiple trees per locus could also be used in order to account for uncertainty), and a network with a given number of reticulations that optimizes the MDC criterion (the lower its value, the more optimal the network) is sought.

As inference based on the MDC criterion does have its limitations in terms of statistical consistency [34] and does not allow for estimating evolutionary parameters beyond the topology of the network, method development turned to statistical inference. To accomplish likelihood-based inference, the multispecies coalescent [7] was extended to the multispecies network coalescent, or MSNC [41, 42]. This extension allowed for deriving the probability distribution of gene trees in the presence of both ILS and reticulation. Heuristic search techniques were developed for maximum likelihood estimation of phylogenetic networks under the MSNC using gene tree estimates as the data [42]. As likelihood calculations on phylogenetic network proved computationally very challenging, a set of techniques was first developed to speed up the exact likelihood calculations [44] and then a pseudo-likelihood measure of phylogenetic networks was devised to allow for faster maximum (pseudo-)likelihood inference [43].

A major challenge facing the aforementioned inference techniques was determining the true number of reticulations, that is, the issue of model complexity. The MDC criterion does not penalize model complexity, and adding more reticulations almost always results in “better” networks under the criterion. Maximum likelihood estimation faces the same issue, and attempts at employing information criteria such as the Akaike Information Criterion (AIC) and Bayesian Information Criterion (BIC) did not provide satisfactory results [42]. We illustrate this issue in Section 4 and discuss how to deal with it in practice when doing either maximum parsimony or maximum likelihood inference. To address the challenge of model complexity more systematically, a Bayesian approach was pursued so that model complexity is accounted for in the prior that is defined on the model. Bayesian inference of phylogenetic networks from gene trees was first devised [37] and then to enable estimation of more parameters and to handle gene tree uncertainty more systematically, methods for Bayesian inference from multiple sequence alignment data and bi-allelic marker data were devised [38, 48]. To speed up computations from bi-allelic markers, a pseudo-likelihood based technique for bi-allelic marker data was developed [47].

Last but not least, as phylogenetic network inference is computationally very expensive (could be orders of magnitude more expensive than tree inference), initial steps have been taken towards scalable phylogenetic network inference using divide-and-conquer techniques [46].

All of the aforementioned inference methods have been implemented in PhyloNet [39], in addition to other features for comparing and analyzing phylogenetic networks [33]. In this paper, we do not review the mathematics or algorithmic techniques behind the methods, as those were reviewed recently [8]; rather, we illustrate the use of these methods on a simulated data set in the hope that the reader will learn how to use the various methods in PhyloNet, how to interpret the results, and what the capabilities and limitations of the methods are. The rest of the paper is organized as follows. In Section 2, we review what a phylogenetic network depicts and how to interpret it. In Section 3, we discuss the nature of the algorithmic techniques employed by PhyloNet for phylogenetic network inference under the various criteria. In Section 4, we illustrate the application of the various inference methods on a simulated data set, and discuss how to interpret the results and how to use the methods with caution, especially with respect to the number of reticulations. Furthermore, we report on the running times of the various analyses in this paper so as to give an idea to the user about the computational requirements of the various inference methods in PhyloNet.In Section 5 we discuss how to analyze large data sets with the current methods in PhyloNet. Given that a set of phylogenetic networks can be inferred, we discuss in Section 6 how to summarize them. In Section 7, we discuss the issue of evolutionary processes underlying the umbrella term *reticulation* and how they cannot be distinguished by methods in PhyloNet (or any method, we believe). We also discuss how PhyloNet can be used to analyze data with polyploids. We end with conclusions in Section 8.

## 2 Reading and Interpreting a Phylogenetic Network

The phylogenetic networks that PhyloNet deals with take the shape of rooted, directed, acyclic graphs (to distinguish them from other types of phylogenetic network [20]). Fig. 1(a) gives an example of a phylogenetic network on seven taxa (e.g., species): C, F, K, L, O, P, and Z. What distinguishes a phylogenetic network from a phylogenetic tree is the presence of *reticulation nodes* in the former. A reticulation node has two parental nodes and its presence denotes that some of the genetic material in the node’s descendants are inherited from one parent while other genetic material in the node’s descendants are inherited from the other parent. For example, the phylogenetic network of Fig. 1(a) depicts a scenario where some loci in P’s genome share a most recent common ancestor with loci from O’s genome, and other loci in P’s genome share a most recent common ancestor with loci from K. Roughly speaking, the percentages of loci belong to these two categories are 35% and 65%, as denoted by the *inheritance probabilities* that label the two *reticulation edges* associated with that reticulation node.^1^

**Figure 1:**
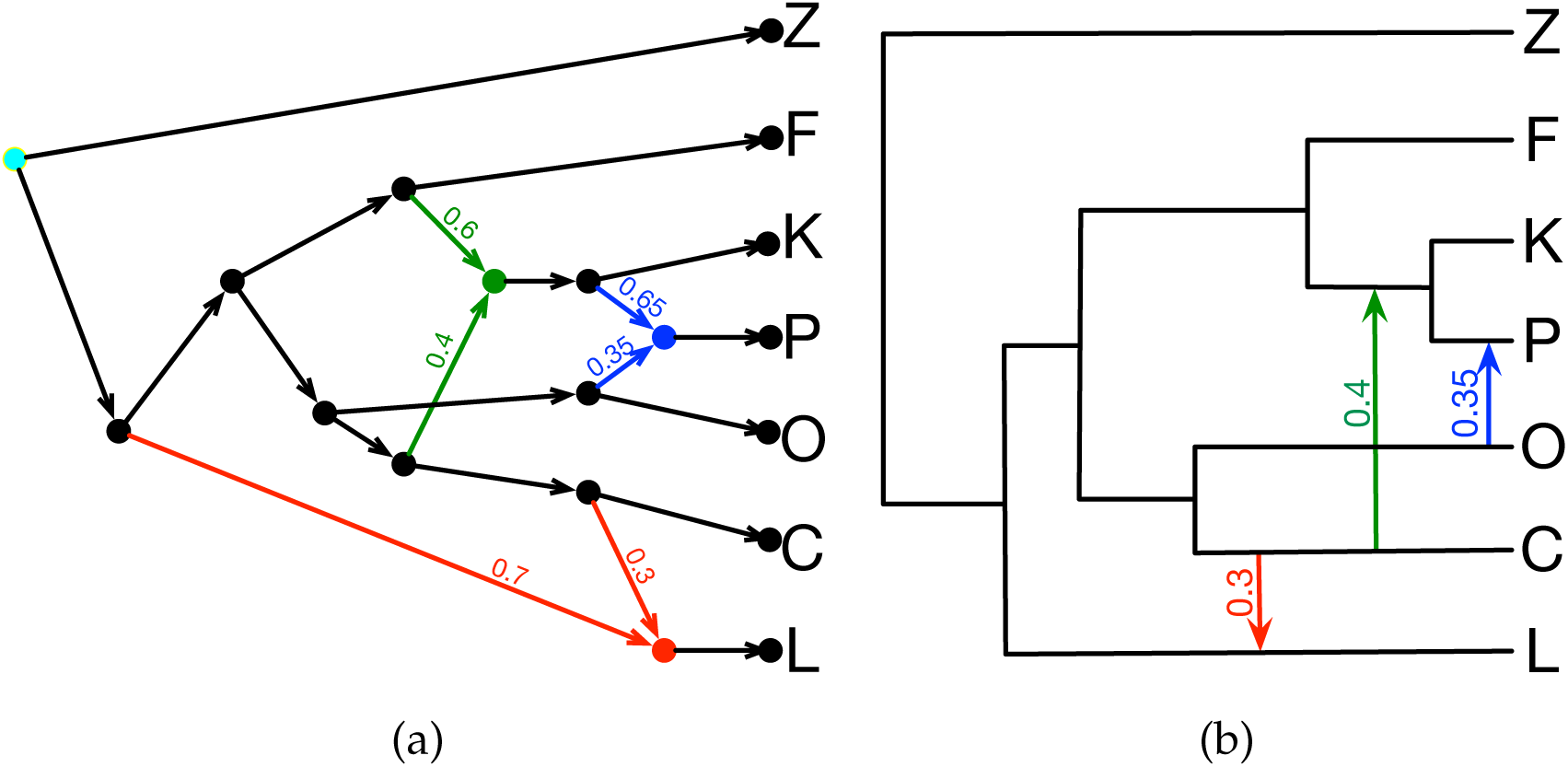
“Reading” a phylogenetic network. (a) The shape, or topology, of a phylogenetic network is a rooted, directed, acyclic graph. The (unique) root is cyan node. The reticulation nodes are colored in red, blue, and green. (b) Depicting the same network in terms of a backbone tree (the solid black lines) and a set of reticulation edges (the red, blue, and green arrows). The inheritance probabilities are shown on the reticulation edges.

For ease of visualization, a phylogenetic network can be drawn as a tree, which we refer to hereafter as “backbone tree,” a set of reticulation edges each of which connects two edges of the backbone tree, as shown in Fig. 1(b). In this case, the user could interpret the backbone tree as the species tree, and the reticulation edges as gene flow events. However, we cannot overemphasize the point that this is only a visualization decision; graph-theoretically, and as far as PhyloNet methods are concerned, the two seemingly different phylogenetic networks in Fig. 1 are exactly the same (more on this issue in Section 7).

A comment about the meaning of a reticulation edge is in order. For actual biological species and populations, a reticulation edge abstracts an epoch of gene flow of unknown duration, but likelihood based species network methods make the simplifying assumption that horizontal gene transfer occurs through instantaneous intermixture. This issue, as well as the performance of Bayesian inference in PhyloNet on data simulated under a model of gene flow are both presented in [38].

### 2.1 Phylogenetic Network Parameters and Their Identifiability

In addition to the inheritance probabilities, the phylogenetic network could have other parameters whose identifiability from inference perspective depends on the criterion employed and data used by the inference method. Three such parameters are branch lengths in coalescent units that are associated with the network’s branches, times that are associated with the nodes (divergence and reticulation times), and population mutation rates that are associated with the branches. Parsimony-based inference using the MDC criterion does not allow for the inference of any of these parameters. Statistical inference using gene tree topologies mainly allows for estimating branch lengths in coalescent units. Statistical inference from the sequence data directly allows for estimating all these parameters (assuming, for example, the generation times are known). However, the identifiability of parameter values depends on the data being used, as we now illustrate. Fig. 2(a) shows a simple 3-taxon phylogenetic network with a single reticulation node where the time associated with the reticulation node is 0.0358. Using the method `MCMC_SEQ` in PhyloNet, which implements Bayesian inference from sequence alignment data [38], the time associated with the reticulation node is unidentifiable when only one individual is sampled from K identifiable, but its estimation improves significantly when two individuals (or alleles) are sampled from species K, as shown in Fig. 2(b) and Fig. 2(c), respectively. The sampled values for the time range between 0 and 0.2 with similar frequencies when a single individual from K is sampled, indicating no confidence in this value. However, when two individuals are sampled, the distribution of the sampled values changes drastically and peaks around the correct value, providing high confidence in the estimated value.

**Figure 2:**
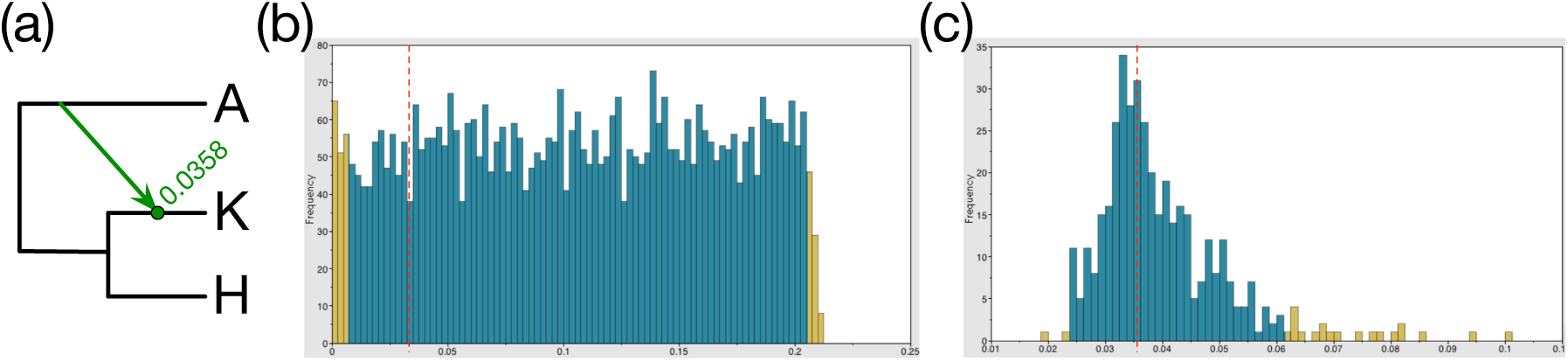
Identifiability of the network’s parameters. (a) A 3-taxon phylogenetic network. Shown in green is the time, in coalescent units, associated with the reticulation node. Histograms of the values sampled for the reticulation node time using `mcmc_seq` when (a) a single allele per taxon is sampled, and (b) when two alleles per taxon are sampled. The dotted vertical lines correspond to the true value (0.0358).

Examples of indistinguishability under the MSNC were discussed in [41], and a more thorough discussion, including with respect to what set of trees characterize a network under the MSNC, was presented in [27, 50, 49].

## 3 Heuristic Searches, Point Estimates, and Posterior Distributions, or, Why Am I Getting Different Networks in Different Runs?

The inference methods listed above and implemented in PhyloNet can be divided into two categories. The first category consists of solutions to optimization problems—problems defined by a solution, or solutions, that optimize a certain criterion. Belonging to this category are `InferNetwork_MP`, `InferNetwork_ML`, `InferNetwork_MPL`, and `MLE_BiMarkers`, which search for networks that optimize the MDC criterion, likelihood and pseudo-likelihood from gene trees and likelihood from bi-allelic markers, respectively. The second category consists of `MCMC_GT`, `MCMC_SEQ`, and `MCMC_BiMarkers` which use Markov chain Monte Carlo simulations to approximate the posterior distribution of the networks and their parameters from gene tree, sequence alignment, and bi-allelic marker data. Methods in the second category do not optimize a criterion.^1^

All problems that these methods are designed to solve are intractable—no exact and practical solutions exist for any of them. By their nature, heuristics and approximations are not guaranteed to solve problems optimally, though well-designed ones perform very well in practice in terms of both accuracy and computational efficiency. When run from different starting points, these methods could return different solutions (e.g., networks) on the same data set. We illustrate why this happens and how to handle the seemingly different results from different runs on an example data set involving trees in Fig. 3.

**Figure 3:**
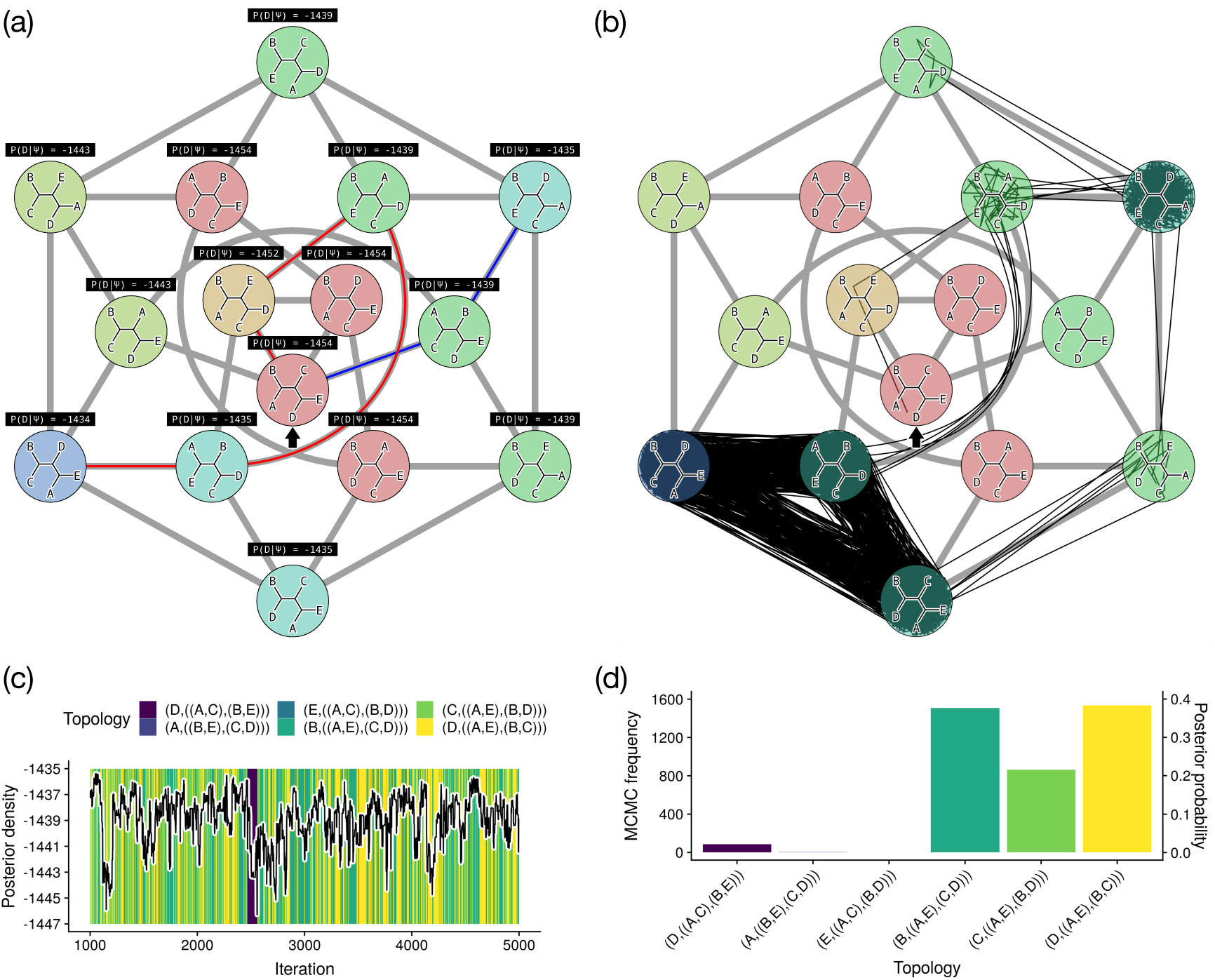
Heuristic search vs. MCMC sampling. (a) Two possible hill climbing paths for 5-taxon unrooted tree topologies using nearest-neighbor interchange (NNI). Phylogenetic likelihoods are for the optimal branch lengths for a given topology. Edges between topologies represent possible NNI moves. The blue path begins from the randomly chosen starting topology (indicated with a solid arrow), and improves the likelihood until a local optimum is reached. The red path begins from the same starting topology but instead by chance improves the likelihood until the global optimum is reached. Topologies are colored in a spectrum spanning red through green to blue from the lowest to highest likelihoods. (b) MCMC for unrooted trees with multiple optima. A 5000-iteration MCMC random walk (black path) is used to estimate the posterior distribution over topologies and branch lengths, with a uniform prior over both, by NNI operators, and by random changes to branch lengths. The walk is initialized from a starting topology (black arrow). Grey lines show changes to tree topologies which are possible using NNI. (c) The posterior density appears well sampled but the minor topology mode is only sampled once during the chain. (d) The number of MCMC samples is exactly proportional to the estimated posterior probability of each topology, a mathematical feature of MCMC.

For the results in the figure, we simulated a sequence alignment of length 300bp, where the first 150bp evolved down the tree (D,((A,C),(B,E))) and the last150bp evolved down the tree (D,((A,E),(B,C))). Starting the search for the maximum likelihood estimate twice from the same starting tree could take the search along two different paths, one that reaches a local optimum that is not the maximum likelihood estimate, while the other does indeed reach the global optimum that is the maximum likelihood estimate (Fig. 3(a)).

In the case of Bayesian MCMC, a random walk starts from a certain tree and traverses the space of trees, not in search of an optimum, but to “collect” information on the posterior distribution over all the possible trees. As Fig 3(b) shows, the random walk visits the different trees, but “circulates” among three topologies more than the other 12 topologies. The posteriors of the trees visited are plotted as in Fig. 3(c), and then the marginal probability distribution on tree topologies is summarized from the samples, as in Fig. 3(d). While one could return the tree (D,((A,E),(B,C))) as the one having the highest marginal probability (the yellow bar is the highest in Fig. 3(d)), the power of Bayesian analysis is that the totality of the results shown in the figure provide more information than just the optimal point. For example, the results in Fig. 3(d) show that the confidence level does not exceed 40% for any of the possible 15 trees. Furthermore, it shows that two trees have comparable probabilities, while a third one is close to them.

Back to network inference, any search strategy begins with one or more starting networks, proposes changes to the network or networks to generate a new candidate or set of candidate networks, and through some evaluation chooses which networks to retain or discard. This is true of the simplest strategy (basic hill climbing), or more complex strategies like stochastic hill climbing, simulated annealing, or genetic algorithms. A starting network may be either randomly generated or it may be a point estimate derived from some other network. A popular way to generate a starting network is using some kind of maximum parsimony algorithm; e.g., `MCMC_GT` uses a starting network with 0 reticulations (i.e., a tree) inferred using the MDC algorithm. The point of using a non-random starting network is to find the optimal network in less time, by beginning the search closer in tree space to the optimum.

Beginning from the starting network, modifications are proposed using “operators” which sample from a distribution of networks that is conditional on the current network; this distribution can be considered the “neighborhood” of the current network. For the MCMC search strategies the mathematical properties of the proposal distribution are important but for point estimate search strategies they are irrelevant; as long as the operators can efficiently traverse the space of networks from the start to the optimum, they are good operators. At some point a stopping condition is reached, and the best network sampled during the search reported as the optimal network. The danger of these search strategies is if a stopping condition causes a search to terminate before the optimal network is sampled, a suboptimal network will be reported as the best network.

How are modifications accepted or rejected? The simplest criterion is to accept modifications which improve the optimality criterion (likelihood or parsimony), and reject those which do not. An algorithm which uses this criterion is called “hill-climbing”, because it attempts to climb up “hills” in the parameter space. A stopping criterion often used for hill climbing is the number of proposals since any proposal was accepted. If say 1,000 modifications are proposed without any being accepted, it is likely that the search has reached *an* optimal point (not necessarily global), and further proposals will also be rejected.

However, just because we can be fairly certain when an optimal point is reached, it is much harder to be certain when the *global* optimum has been reached. In the space of networks (or other parameters) there may be several “hills” of good scores (either high likelihoods, posterior densities, or small parsimony scores). Because basic hill climbing will not climb down a hill, if the current network is sampled from one of the *local* optima far from the global optimum, the global optimum may be beyond the neighborhood of the current state and the search will get stuck on top of the hill (as illustrated in the example of Fig. 3).

If a hill climbing algorithm is run multiple times with different starting seeds, a different series of proposals will be made for each run. While one or more runs may be stuck in local optima, hopefully at least one will by chance reach the global optimum and report it as the best network. Therefore when using a simple hill climbing method, users must execute it multiple times with different starting seeds. The network with the best score across all these runs should be reported as the best point estimate.

## 4 Illustrating the Various Inference Methods in PhyloNet

For the network Fig. 1, we generated 100 gene trees with two individuals per species using the program `ms` [12]. We set the population mutation rate at 0.02, base frequencies of A, C, G and T at 0.2112, 0.2888, 0.2896, and 0.2104, respectively, and the transition probabilities at [0.2173, 0.9798, 0.2575, 0.1038, 1, 0.2070].

We removed the outgroup Z and ran the phylogenetic network inference methods `InferNetwork_MP`, `InferNetwork_ML`, `InferNetwork_MPL` and `MCMC_GT` based on true gene trees and inferred gene trees from IQTREE [23]. In terms of the accuracy of gene tree estimates, 72 out of 100 gene trees inferred by IQTREE are topologically identical to true gene trees. The average error of gene tree branch lengths, which is computed as the rooted branch score distance [14] between each pair of inferred and true gene trees normalized by the branch length of the true gene tree, is 6.58%.

For each inference method, the maximum number of reticulations is set to be 0, 1, 2, 3 and 4, respectively. The number of processors used is 8, and all other parameters are the default settings.

We refer to the tree drawn with black lines (the network without the three arrows) in Fig. 1 as the backbone tree.

### 4.1 Inference Under the MDC Criterion

Phylogenetic networks inferred under the MDC criterion (the `InferNetwork_MP` command) from the true gene tree topologies are shown in Fig. 4. As the figure shows, the optimal tree (i.e., network with no reticulation nodes) inferred is the backbone tree of the true network. Furthermore, as the number of reticulations allowed was increased, the three reticulation events were identified one by one, until the true network was inferred (Fig. 4(d)).

**Figure 4:**
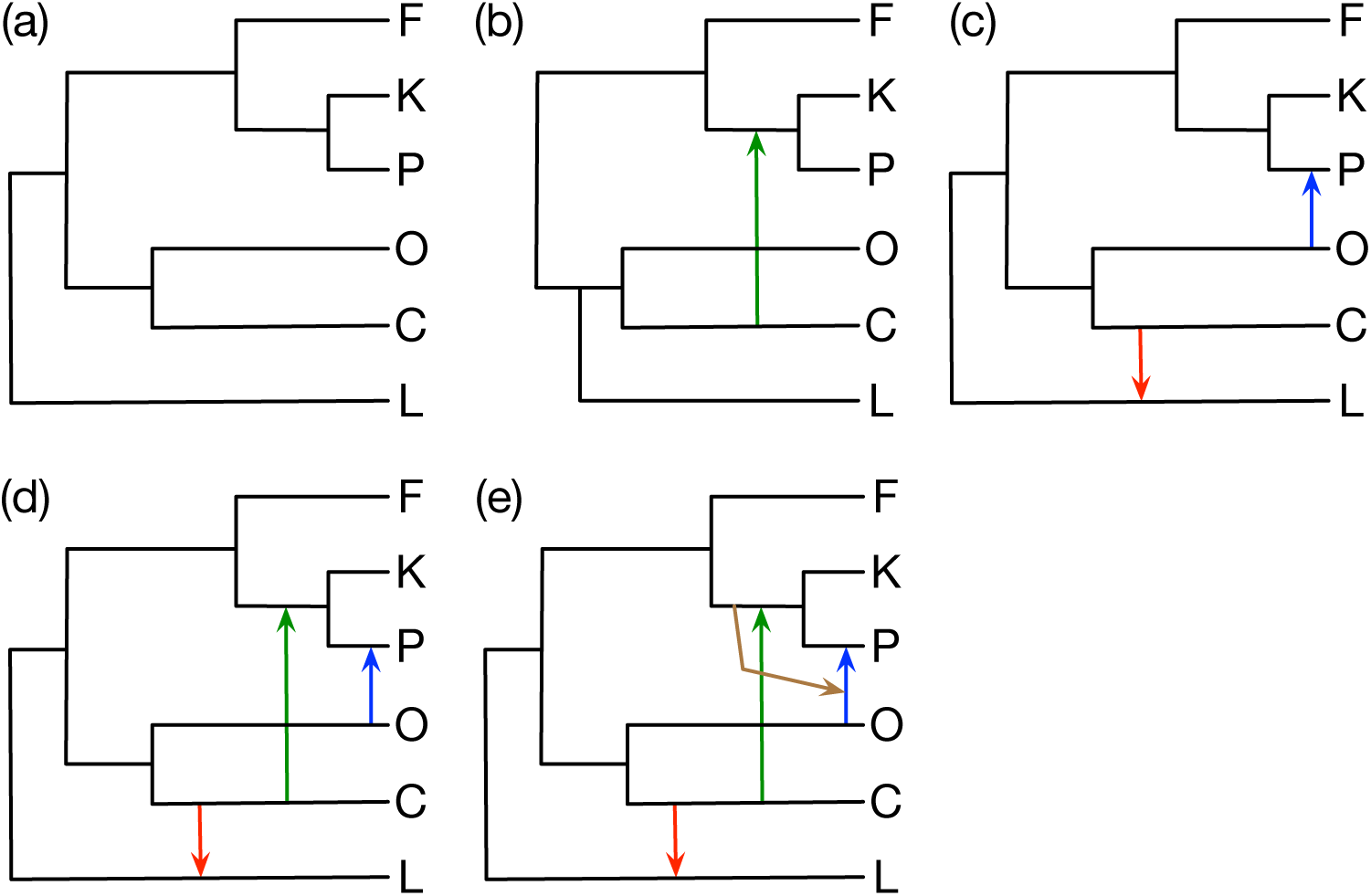
Inference results using the MDC criterion on the true gene tree topoloies. (a)-(e) The optimal networks inferred when the maximum number of reticulations was set to 0, 1, 2, 3, and 4, respectively. The MDC scores of the five networks in (a)-(e) are 363, 276, 175, 67 and 63, respectively.

Three points are worth highlighting based on the results of Fig. 4. First, while the reticulation between C and (K,P) was correctly inferred when the maximum number of allowed reticulations was set at 1, the placement of L is incorrect (Fig. 4(b)). There could be two explanations for this error in the network. One is that the network shown has a better score under the MDC criterion than one with the correct placement of L. The second is that the network with the correct placement of L has a better score under the MDC criterion, but that the heuristic for inferring the optimal network with a single reticulation was not run long enough to find it. To answer this question, we compared the MDC score of the network in Fig. 4(b) and the MDC score of the network that differs only in correcting the placement of L so that it matches that in the true network. We found that the network (a) with the reticulation from C to (K,P) has MDC score 278, which means the network with the wrong placement of L has a better score under MDC. However, the difference in the MDC score between the two networks is only 2, an indication that the MDC criterion, in this case, cannot place L with high confidence. It is important to note here that this type of analysis cannot be done with real data, as the true network is unknown. However, if the biologist has an evolutionary hypothesis in mind (e.g., some modification to the inferred network), they can compute the MDC score of that network and compare it to the one inferred by the method (the `DeepCoalCount_network` command).

Second, observe that the network with two reticulations (Fig. 4(c)) is not simply the network with one reticulation (Fig. 4(b)) plus an additional reticulation. In fact, the two networks are disjoint in terms of the reticulation events they model. This observation illustrates the important point that under the MDC criterion, an optimal network with *k* + 1 reticulations cannot necessarily be obtained by first inferring an optimal network with *k* reticulations and then searching for one additional reticulation to add to it.

Third, while the true number of reticulations is three, the method inferred a network with four reticulations when the maximum number of reticulations allowable was set at four. This illustrates that the MDC criterion by itself is not sufficient to determine the true number of reticulations. One technique to use in practice, yet is not foolproof under all conditions, is to inspect the improvement to the MDC score as more reticulations are inferred. In this case, as the number of reticulations identified went from 0 to 4, the MDC score dropped according to the sequence 363, 276, 175, 67, and 63, respectively. Translating these MDC scores into subsequent rates of change (subtracting two consecutive numbers and dividing by the larger) yields improvements in the MDC scores of roughly 24%, 37%, 62%, and 6%. Notice that the improvement in the MDC score from the network with three reticulations to the one with four reticulations is much smaller than the other three improvements, a good indication that three reticulations is the correct number in this case. As the MDC score decreases with the addition of reticulations and to avoid artificial percentages due to small denominators^1^, one can also inspect the rate of change with respect to the MDC score of the tree: (363 −276) /363 ≈ 24%, (363 −175) /363 ≈ 52%, (363− 67) /363 ≈ 82%, and (363− 63) /363 ≈ 83%. This inspection makes it even clearer that the fourth added reticulation is very unlikely to be a real one but rather a reflection of the fact that the MDC score cannot get worse by adding more reticulations to the same network.

As gene trees are estimated from data in practice, we repeated the analysis but with the data consisting of gene tree topologies estimated using IQTREE, and rooted at outgroup Z. The results are shown in Fig. 5. The main observations here are that the method inferred the correct network with three reticulations, despite the error in the gene tree estimates, and that the order in which the reticulation events were inferred is not consistent with that in the case of true gene trees. The placement of L is now wrong when the method inferred a network with two reticulations, and the fourth “fake” reticulation is different from the one identified before. However, there are no generalizable patterns here. All these observations could be affected by the quality of the estimated gene trees.

**Figure 5:**
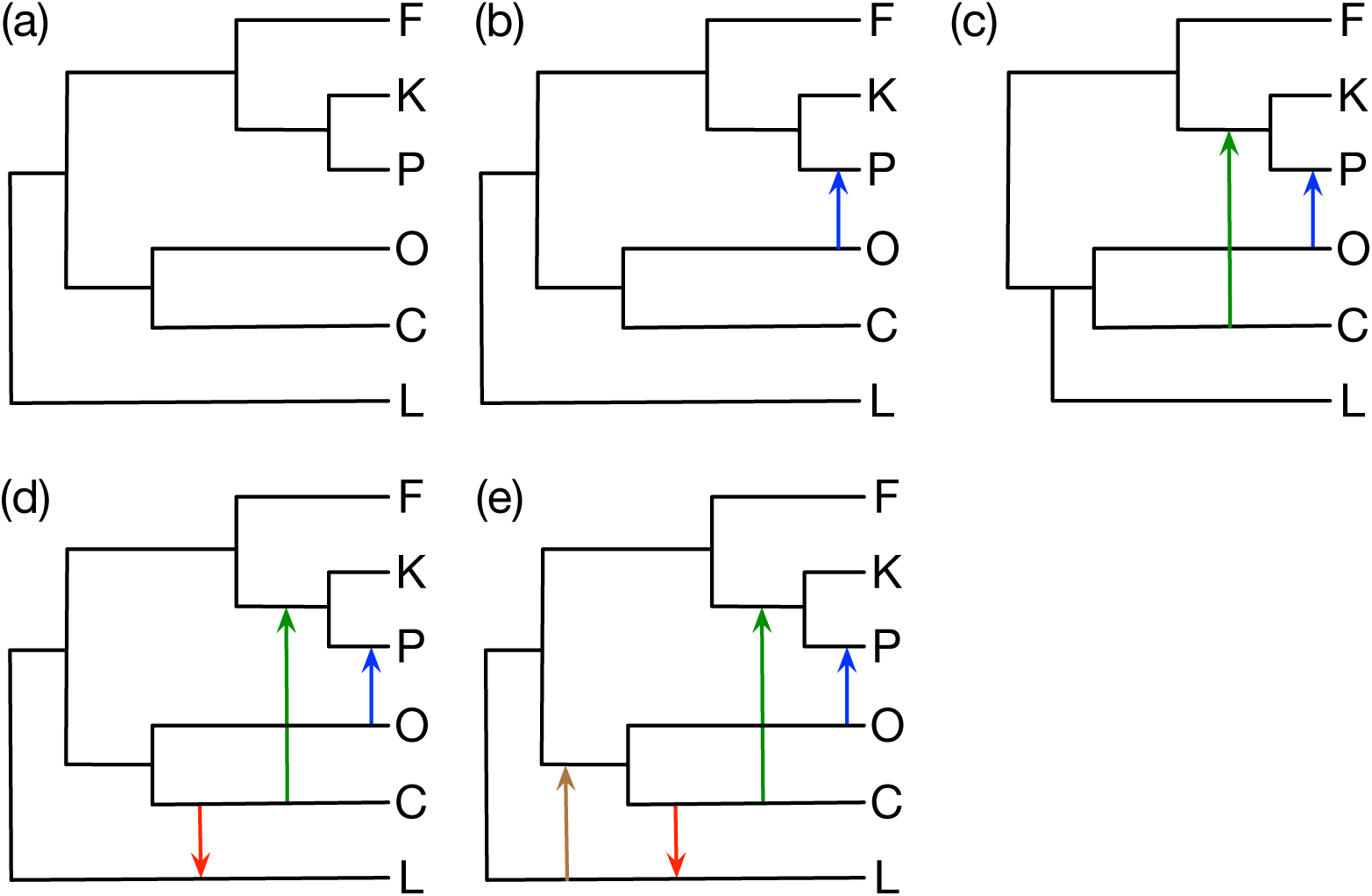
Inference results using the MDC criterion on the gene tree topologies estimated by IQTREE. (a)-(e) The optimal networks inferred when the maximum number of reticulations was set to 0, 1, 2, 3, and 4, respectively. The MDC scores of the five networks are 384, 270, 175, 100, and 88, respectively.

The improvements to the MDC scores are now (384− 270)/ 384≈30%, (384−175)/ 384 ≈ 54%, and (384 −100)/ 384 ≈74%, (384 − 88)/ 384 ≈ 77%. We observe that in this case as well the criterion points to three as the correct number of reticulations.

Having illustrated MDC, especially that it worked perfectly on this data set, it is important to note that inference under the MDC criterion is not statistically consistent in the case of species trees [34]. Therefore, there would be cases where the wrong network is inferred under the MDC criterion. However, we have observed that the criterion does very well in general [40].

### 4.2 Maximum Likelihood Inference

Phylogenetic networks inferred under maximum likelihood (the `InferNetwork_ML` command) from the gene tree topologies are shown in Fig. 6.

**Figure 6:**
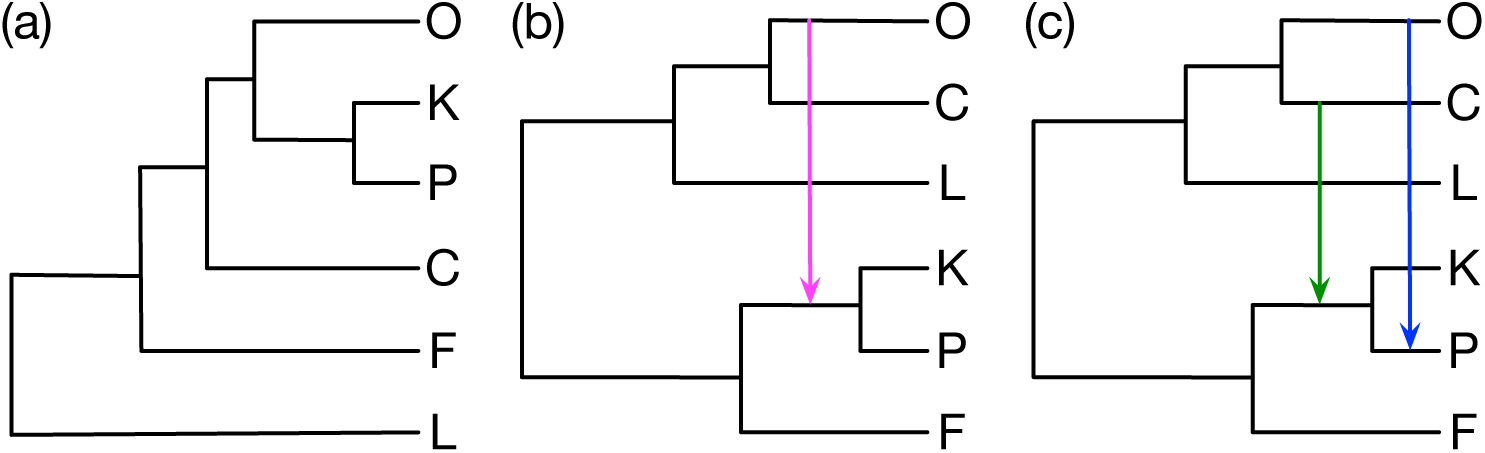
Maximum likelihood inference results on gene tree topologies. (a)-(c) The optimal networks inferred when the maximum number of reticulations was set to 0, 1, and 2, respectively. While the same network topologies were inferred from the true gene trees as well as from the gene trees estimated by IQTREE, the network parameters, as well as their likelihoods, differed. The log likelihoods of the three networks in (a)-(c) when using the true gene trees are -790, -686, and -653, respectively. The log likelihoods of the three networks in (a)-(c) when using the IQTREE gene tree estimates are -792, -719, and -635, respectively. ML inference with maximum number of reticulations set at 3 and 4 did not finish within 24 hours. The log likelihood of the true network is -388 for true gene trees and -624 for IQTREE.

Given the likelihood calculations are very expensive when gene tree topologies are used as input [49, 47, 8], analyses where the maximum number of reticulations was greater than two were infeasible within the time allotted. For analyses where the number of inferred reticulations was capped at two or fewer, inference based on the true and estimated gene tree topologies resulted in the same networks, which is a reflection of the method’s robustness to gene tree error (up to a certain level).

The MDC criterion is not a multispecies coalescent criterion per se; rather, it is a simple criterion that aims at minimizing conflict between the input gene trees and the species phylogeny. On the other hand, maximum likelihood inference in this case *explicit* makes use of the multispecies (network) coalescent. The results are shown in Fig. 6, and even more so in Fig. 7 where branch lengths of the gene trees are used, highlight the big impact of gene flow on species *tree* inference, an issue that has been discussed and analyzed before [49, 38, 8, 1].

**Figure 7:**
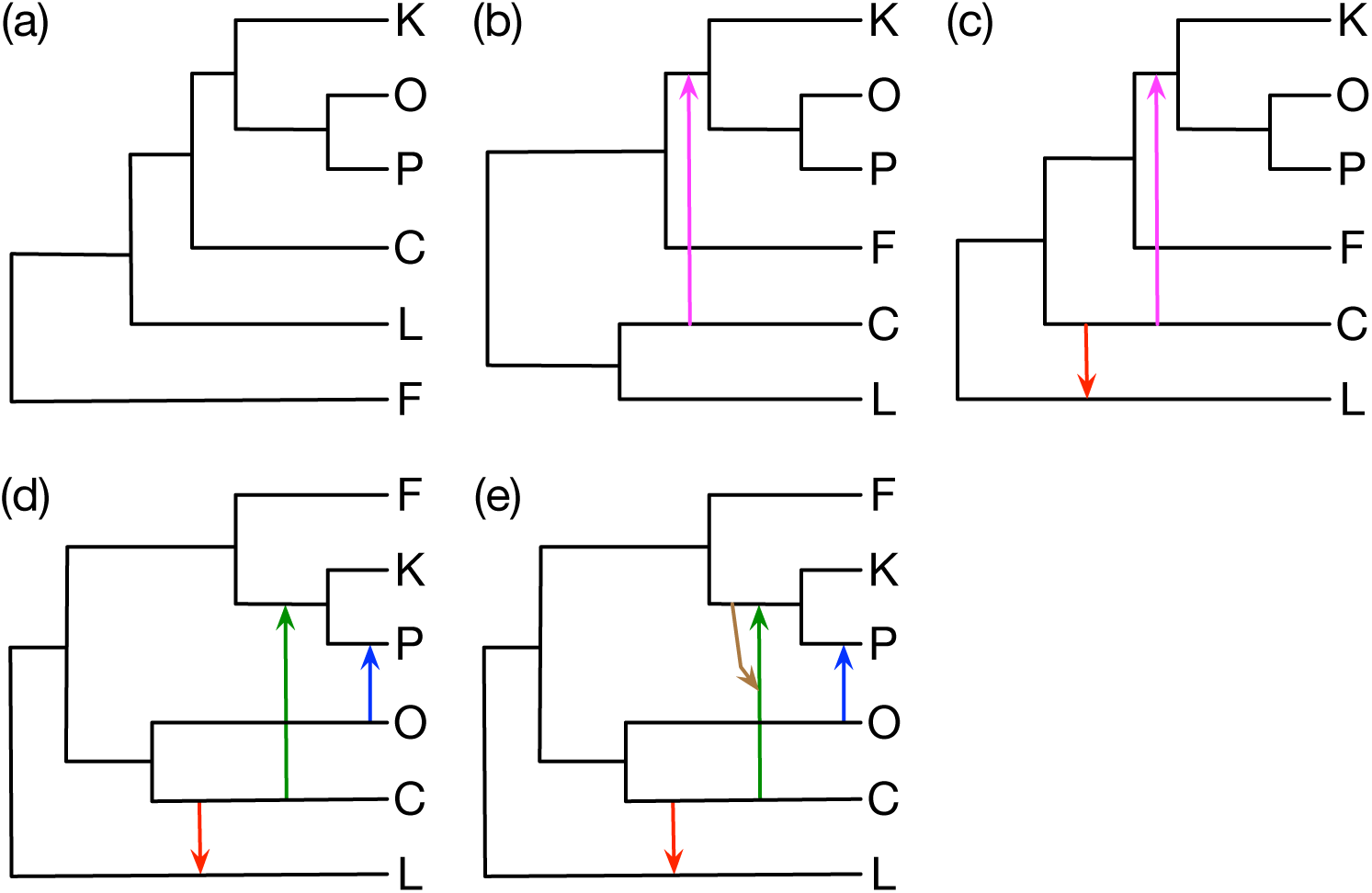
Maximum likelihood inference results on the true gene trees (topologies and branch lengths). (a)-(e) The optimal networks inferred when the maximum number of reticulations was set to 0, 1, 2, 3, and 4, respectively. The log likelihoods of the five networks in (a)-(e) are -11925, -6292, -3492, -1642, and -1494, respectively. The log likelihood of the true network is -1408.

When a tree is inferred under ML (that is, the maximum reticulation is set at 0), the tree (Fig. 6(a)) does not match the true backbone tree. The reason for this is that under the multispecies coalescent, to explain in a tree model the clades that group O and P genes together due to gene flow (the blue arrow in the true network), O is grouped with P and K. Similarly, due to gene flow between C and (K,P) (the green arrow in the true network), C is grouped with the clade the contains P and K before F is. This illustrates the issues that could arise when a species tree is inferred despite the fact that gene flow had occurred, and is congruent with previous results showing that the species tree methods ASTRAL and NJst infer erroneous clades under ILS and gene flow [31]. As reticulations are allowed during inference, the clades, as well as the true reticulations, start to get recovered correctly, as shown in Fig. 6(c).

As we mentioned, this issue is even more pronounced when the ML inference utilizes the branch lengths (effectively the coalescent times) in the gene trees. As Fig. 7(a) shows, the inferred tree, while very different from the true backbone tree, makes much sense when the information on coalescent times is taken into consideration. The earliest possible (looking backward in time from the leaves toward the root) coalescent event can happen between alleles from O and P due to gene flow between the two species (the blue arrow in the true network). The second earliest possible coalescent event can happen between alleles from P and K, and so on. These temporal constraints defined by the phylogenetic network and the fact that a coalescence between two alleles from two species cannot occur before the divergence of these two species from their MRCA or the most recent episode of gene flow between them define a partial order as illustrated in Fig. 8. This partial order explains the species tree inferred and shown in Fig. 7(a). However, as the permitted number of reticulations is increased, the tree and network structures are fully and correctly inferred, as shown in Fig. 7(d). As with inference based on the MDC criterion, maximum likelihood without a penalty for model complexity cannot determine the number of reticulations, as illustrated in the network of Fig. 7(e).

**Figure 8:**
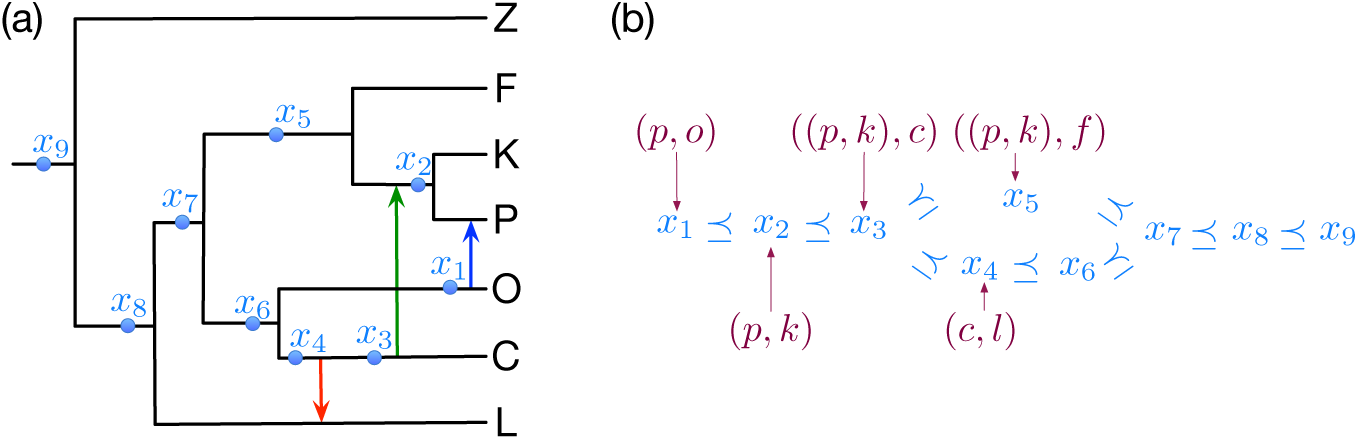
The constraints that coalescent times impose when a species tree is inferred in the presence of gene flow. (a) Coalescent events can occur at any internal branch in the phylogenetic networks, including the points *x*_1_, …, *x*_9_. The backbone tree defines a partial order on those nine points: *x*_1_⪯ *x*_6_, *x*_2_ ⪯ *x*_5_, *x*_3_ ⪯*x*_4_, *x*_4_ ⪯ *x*_6_, *x*_5_ ⪯ *x*_7_, *x*_6_ ⪯ *x*_7_, *x*_7_ ⪯ *x*_8_, and *x*_8_ ⪯ *x*_9_. The three reticulation events impose additional order. For example, due to gene flow, the smallest possible time at which *x*_1_ could occur is smaller than or equal to the smallest possible time at which *x*_2_ could occur, due to alleles from P and O coalescing before alleles from P and K coalesce, thus adding *x*_1_⪯ *x*_2_. (b) The partial order defined by the network and smallest possible coalescent times of *x*_1_, …, *x*_9_.

Finally, results based on gene trees with estimated topologies and branch lengths are similar to results based on true gene tree topologies and branch lengths, as shown in Fig. 9. However, while the method infers the true phylogenetic network topology with three reticulations, the branch lengths are different from those of the true one. In fact, the likelihood of the true network (true topology and true branch lengths) given the estimated gene trees is 0, the reason being that the coalescent times are underestimated for some of the gene trees forcing the network to an nonviable model (e.g., species A and B diverged from their MRCA at time *t*, and alleles from A and B coalesce at time *τ*; having *τ* < *T* is not a viable setting). The issue of underestimating the coalescent times in the gene trees, as well as the implications thereof on species tree estimation, was discussed in [6]. Unless one is certain of the gene tree branch lengths or coalescent times, we recommend against inference that utilizes gene tree branch lengths, despite the fact that it is computationally much less demanding than the inference based on gene tree topologies alone.

**Figure 9:**
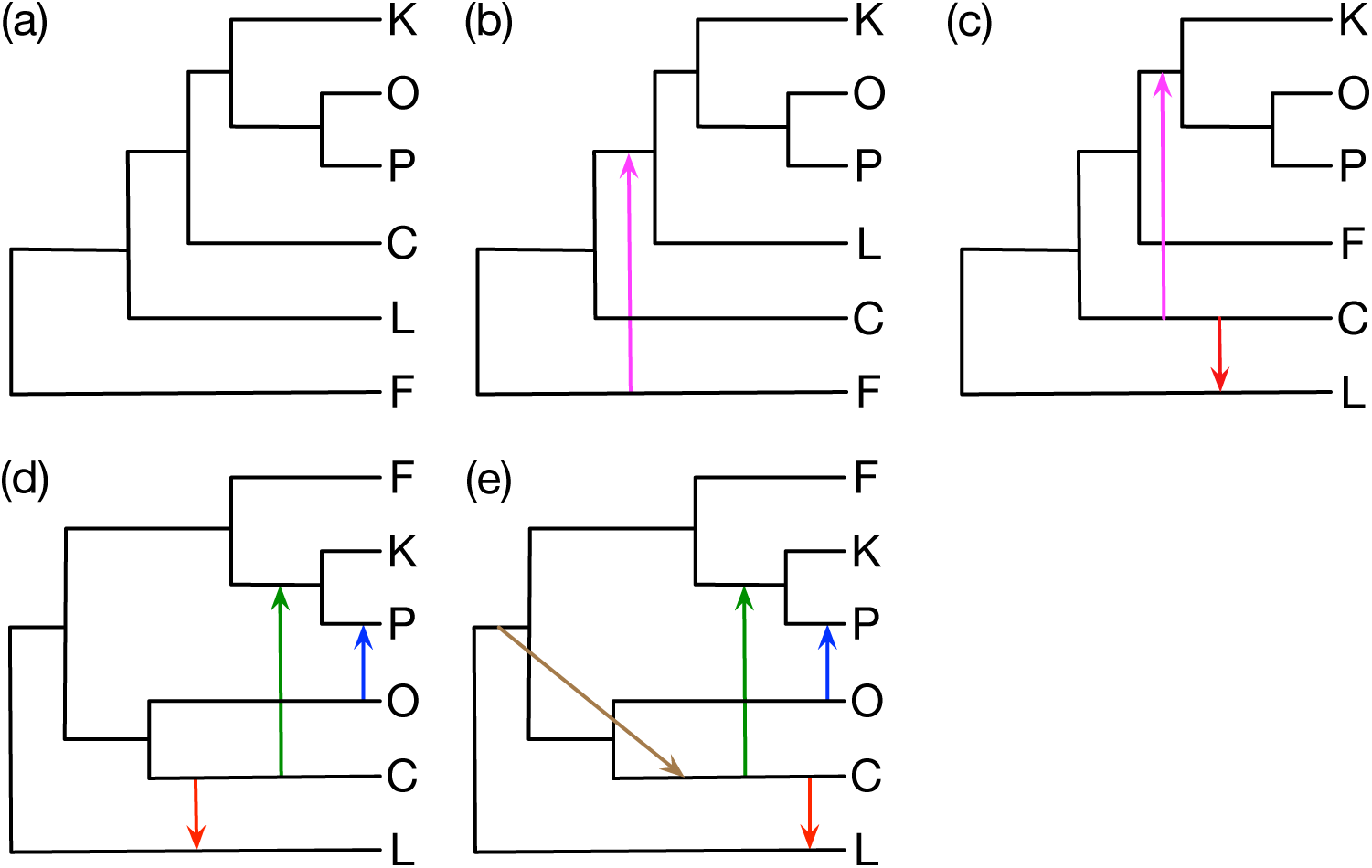
Maximum likelihood inference results on the gene trees (topologies and branch lengths) estimated by IQTREE. (a)-(e) The optimal networks inferred when the maximum number of reticulations was set to 0, 1, 2, 3, and 4, respectively. The log likelihoods of the five networks in (a)-(e) are -15683, -9232, -5977, -4044, and -3653, respectively. The true network (true topology and branch lengths) is not viable given the gene tree estimates (see text).

### 4.3 Maximum Pseudo-likelihood Inference

Phylogenetic networks inferred under maximum pseudo-likelihood (the `InferNetwork_MPL` command) from the true and inferred gene tree topologies are shown in Fig. 10 and Fig. 11, respectively.

**Figure 10:**
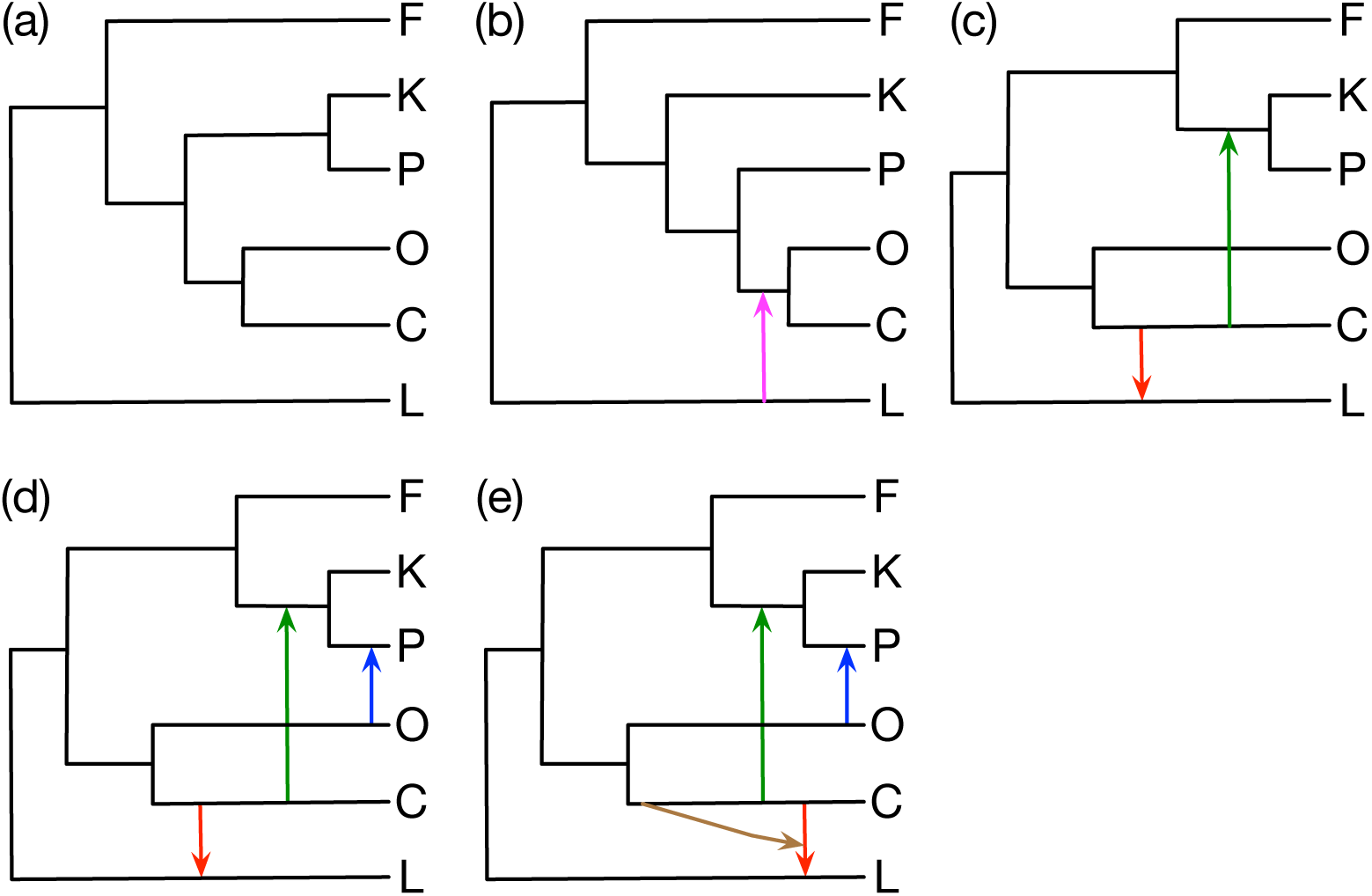
Maximum pseudo-likelihood inference results on the true gene tree topologies. (a)-(e) The optimal networks inferred when the maximum number of reticulations was set to 0, 1, 2, 3, and 4, respectively. The log pseudo-likelihoods of the five networks in (a)-(e) are -13449, -11992, -10791, -10314, and -10311, respectively. The log pseudo-likelihood of the true network is -10341.

**Figure 11:**
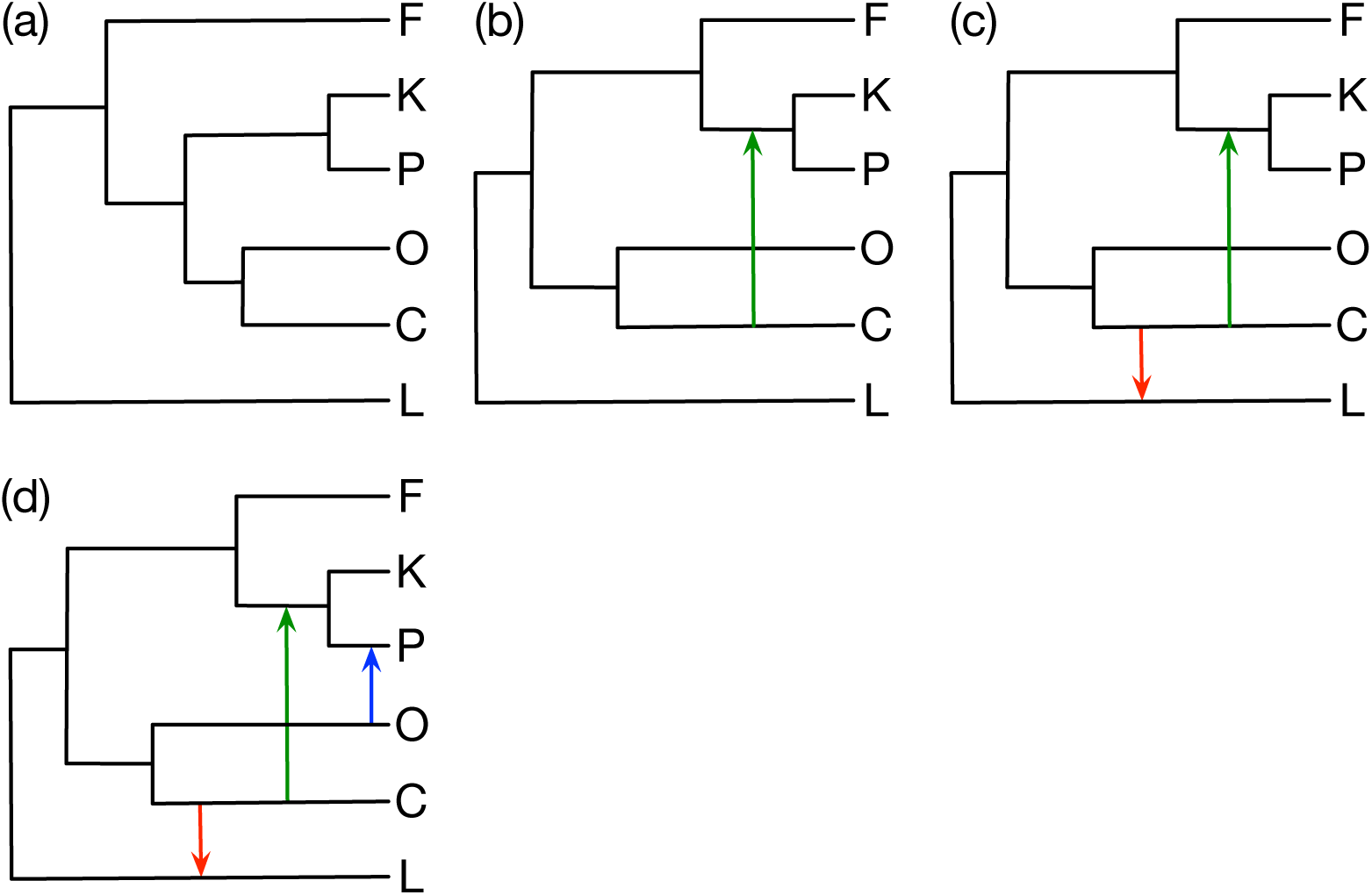
Maximum pseudo-likelihood inference results on the gene tree topologies estimated by IQTREE. (a)-(c) The optimal networks inferred when the maximum number of reticulations was set to 0, 1, and 2, respectively. (d) The optimal network when the maximum number of reticulations was set to 3 and to 4. The log pseudo-likelihoods of the four networks in (a)-(d) are -14582, -13339, -12540, and -12079, respectively. The log pseudo-likelihood of the true network is -13707.

As the pseudo-likelihood of a network is efficiently computable, analyses now were run with up to four reticulations allowed. Using this method we obtained more accurate results than maximum likelihood, particularly in the case of an inferred species tree where maximum pseudo-likelihood seems to be robust the presence of gene flow. Furthermore, it appears easier to identify the correct number of reticulations using pseudo-likelihood. When estimated gene trees were used, the pseudo-likelihood for the best network with three reticulations was identical to that with four reticulations. When the true gene trees were used, the log pseudo-likelihood of the best networks with three and four reticulations found differed by only 3.

We also inferred phylogenetic networks under maximum pseudo-likelihood from bi-allelic markers (the `MLE_BiMarkers` command), and the results are shown in Fig. 12. Except for the species tree in Fig. 12(a) and the incorrect placement of L in Fig. 12(c), the results are very good, with the true network inferred when the maximum number of reticulations is set to 3. It is worth mentioning that when the inference was run with the maximum number of reticulations set to 4, the search did not find any network with better pseudo-likelihood than the one with 3 reticulations shown in Fig. Fig. 12(d). This is almost the same pattern we observed above with maximum pseudo-likelihood using gene tree topologies where the network with four reticulations has almost the same pseudo-likelihood as the one with three reticulations.

**Figure 12:**
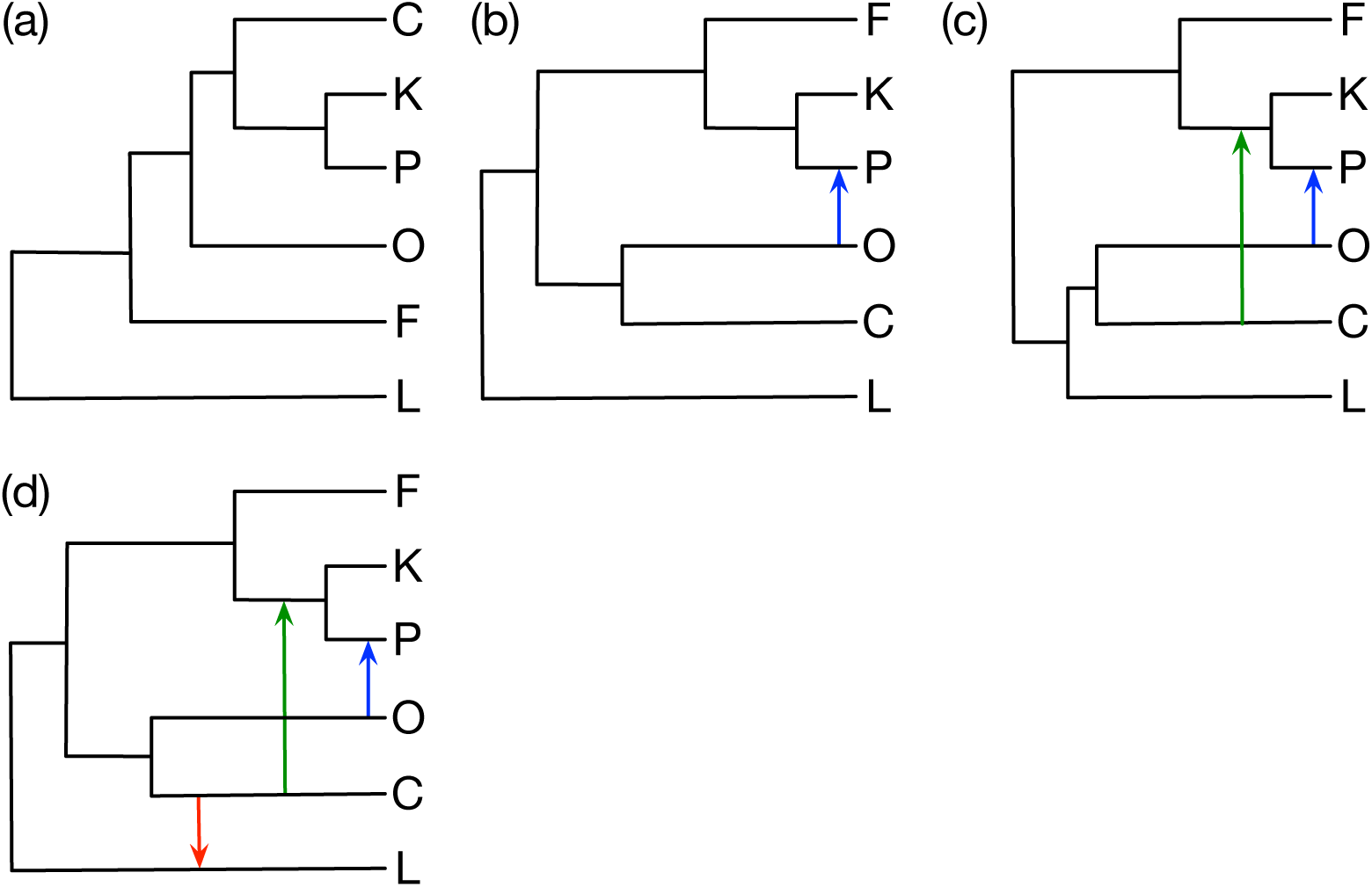
Maximum pseudo-likelihood inference results using bi-allelic marker data. (a)-(d) The optimal networks inferred when the maximum number of reticulations was set to 0, 1, 2, and 3, respectively. The log pseudo-likelihoods of the five networks in (a)-(d) are -677309, -676093, -675271, and -675165, respectively.

### 4.4 Bayesian Inference

PhyloNet has three functionalities that allow for Bayesian inference of phylogenetic networks: `MCMC_GT` infers phylogenetic networks using Bayesian MCMC from gene tree topologies [37], `MCMC_BiMarkers` works directly on bi-allelic marker data [48], and `MCMC_SEQ` infers phylogenetic networks from sequence alignment data [38]. The former two are computationally very demanding and would not converge within 24 hours on the simulated data set used here. For `MCMC_GT`, the computational demands are similar to those of `InferNetwork_ML` when gene tree topologies alone are used. For `MCMC_BiMarkers`, the analytical computation of the integration over all gene trees becomes infeasible for certain phylogenetic networks, as illustrated in [47, 8]. Therefore, we analyzed the data set using `MCMC_SEQ` which, unlike the other two, does not compute the joint integration of species network and gene tree histories entirely analytically; instead, it samples gene trees with their coalescent times and calculates the density function of gene trees. The results are shown in Fig. 13.

**Figure 13:**
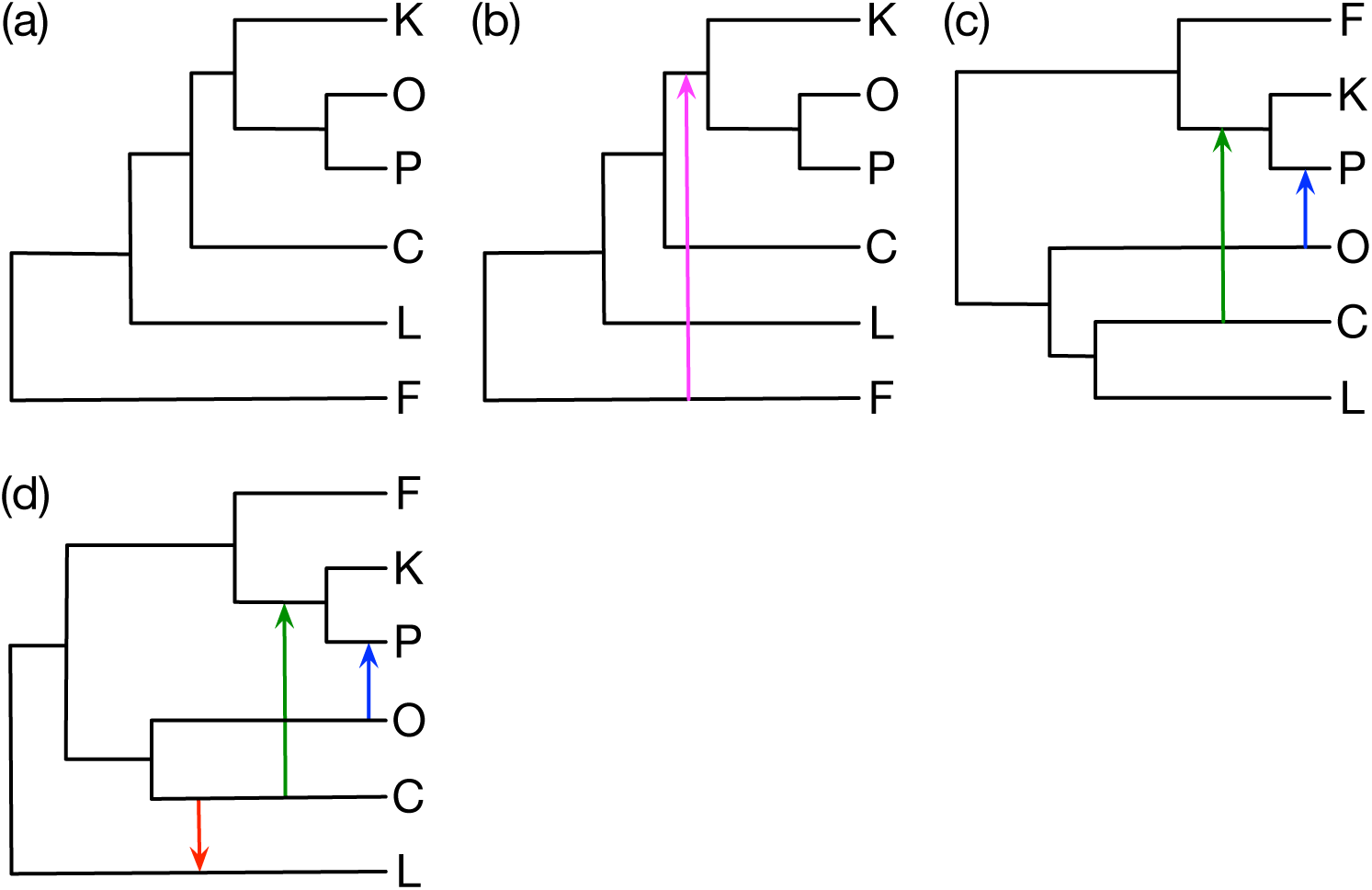
Bayesian inference results on the sequence alignment data. (a) The tree the appeared in 99.7% of the collected samples when the maximum number of reticulations was set to 0. (b) The network that appeared in 100% of the collected samples when the maximum number of reticulations was set to 1. (c) The network that appeared in 100% of the collected samples when the maximum number of reticulations was set to 2. (d) The network that appeared in 100% of the collected samples when the maximum number of reticulations was set to 3 or to 4.

We make three observations. First, similar to the results based on maximum likelihood when the true gene trees with their branch lengths are used (Fig. 7), the species tree (when the analysis is constrained as to not allow any reticulations) has incorrect groupings of the taxa. The explanation for this is the same as that illustrated above with Fig. 8. Second, as the number of reticulations allowed is increased, the inference converges onto the correct network. Third, even when the maximum number of reticulations allowed was set at 4, the method inferred the network with the correct number of reticulations (3), demonstrating the ability of Bayesian inference to determine the correct number of reticulations, by imposing a prior distribution on the number of reticulations.

### 4.5 Running Time

While phylogenetic analyses take non-trival amounts of computational time to perform, generally it will take up the minority of time required for a study which requires fieldwork, sample preparation, and so on. Previous work has shown that Bayesian inference of species trees from sequences scales with the size of the data set employed roughly following a power law [25]. The CPU time required to achieve a posterior density ESS of 200 on our test data set is consistent with this trend for a given number of individuals per species, when inferring a species tree by setting the maximum number of reticulations to zero (Fig. 14).

**Figure 14:**
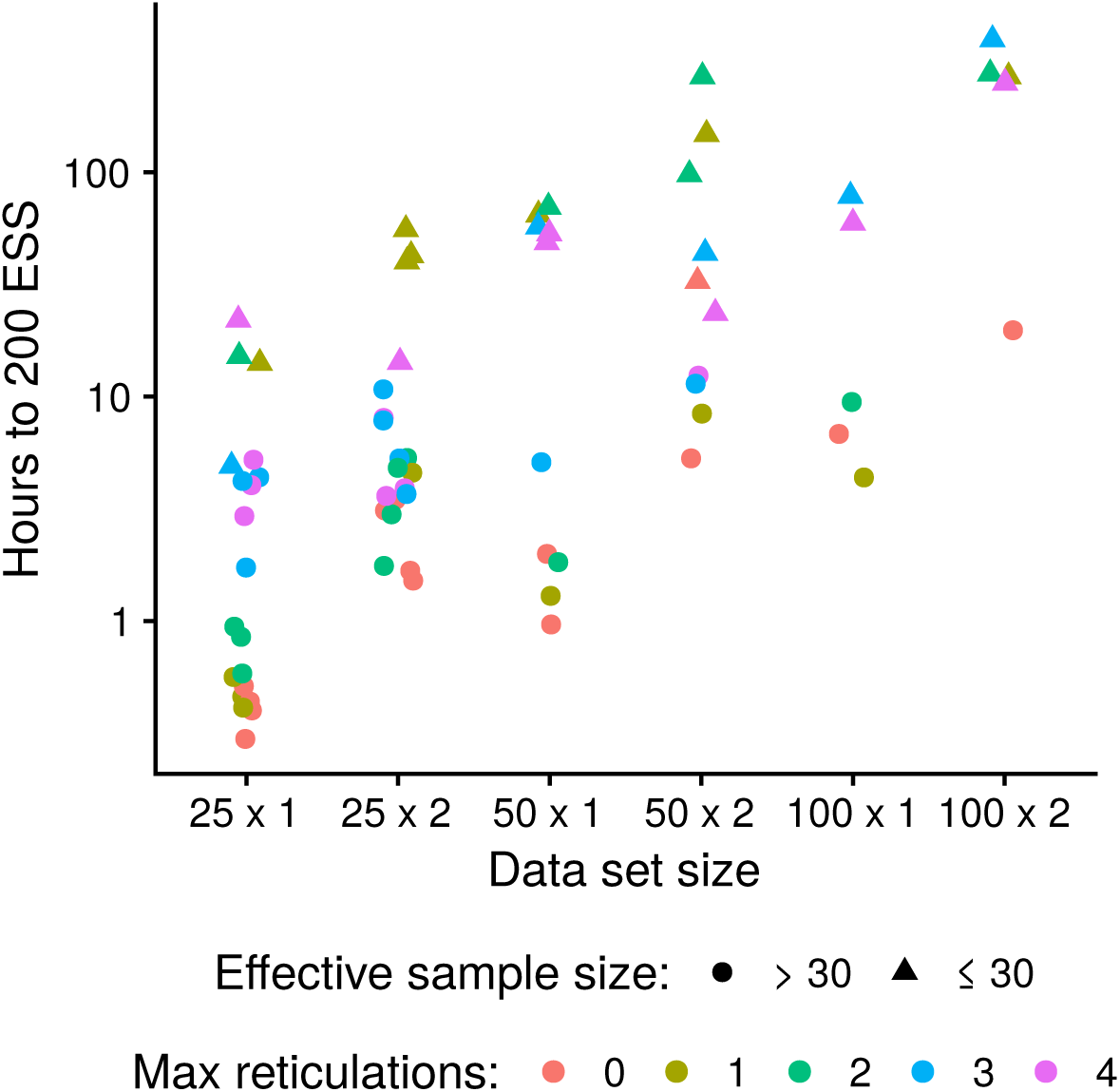
Running time requirements of MCMC_SEQ for the analysis reported in Section 4.4. Two chains were run for each combination of data set and maximum reticulations, then concatenated after removing 50% burn-in before computing effective sample size (ESS). The time to a posterior density ESS of 200 was extrapolated from the ESS of the concatenated chains. CPU time was halved to account for the time taken on the sampled removed as burn-in. The full 100 locus data set was segmented into four 25 locus and two 50 locus data sets. Data set size is given as number of loci by number of individuals per species. Max reticulations refers to the maximum number of reticulations permitted.

Low ESS values may be the result of a chain which is still in its burn-in phase, or a chain which is stuck in a local optima, or a chain which has reached the mode of the stationary distribution but has a high autocorrelation time. After concatenating the posterior samples from two independent chains for each combination of data set and maximum number of reticulations, chains which are had not finished burning in or were stuck in different local optima will cause ESS to be very low (we used less than 30 as a rule-of-thumb). The calculated time to 200 ESS, extrapolated from these very small ESS values, will not be reliable.

Judging from the more reliable extrapolated times, increasing either the maximum number of reticulations permitted, or the number of individuals per species, results increases required CPU time. The time required for the 50 × 1 (loci × individuals per species) analyses were similar to the 25 × 2 analyses, and the 100 × 1 times were similar to 50 × 2, suggesting the total number of sequences in a data set is the key variable determining required time (Fig. 14).

Table 1 shows the running times of the various network inferences we discussed above on NOTS (Night Owls Time-Sharing Service), which is a batch scheduled High-Throughput Computing (HTC) cluster. If the inference exceeded 192 CPU hours, it was terminated.

**Table 1:**
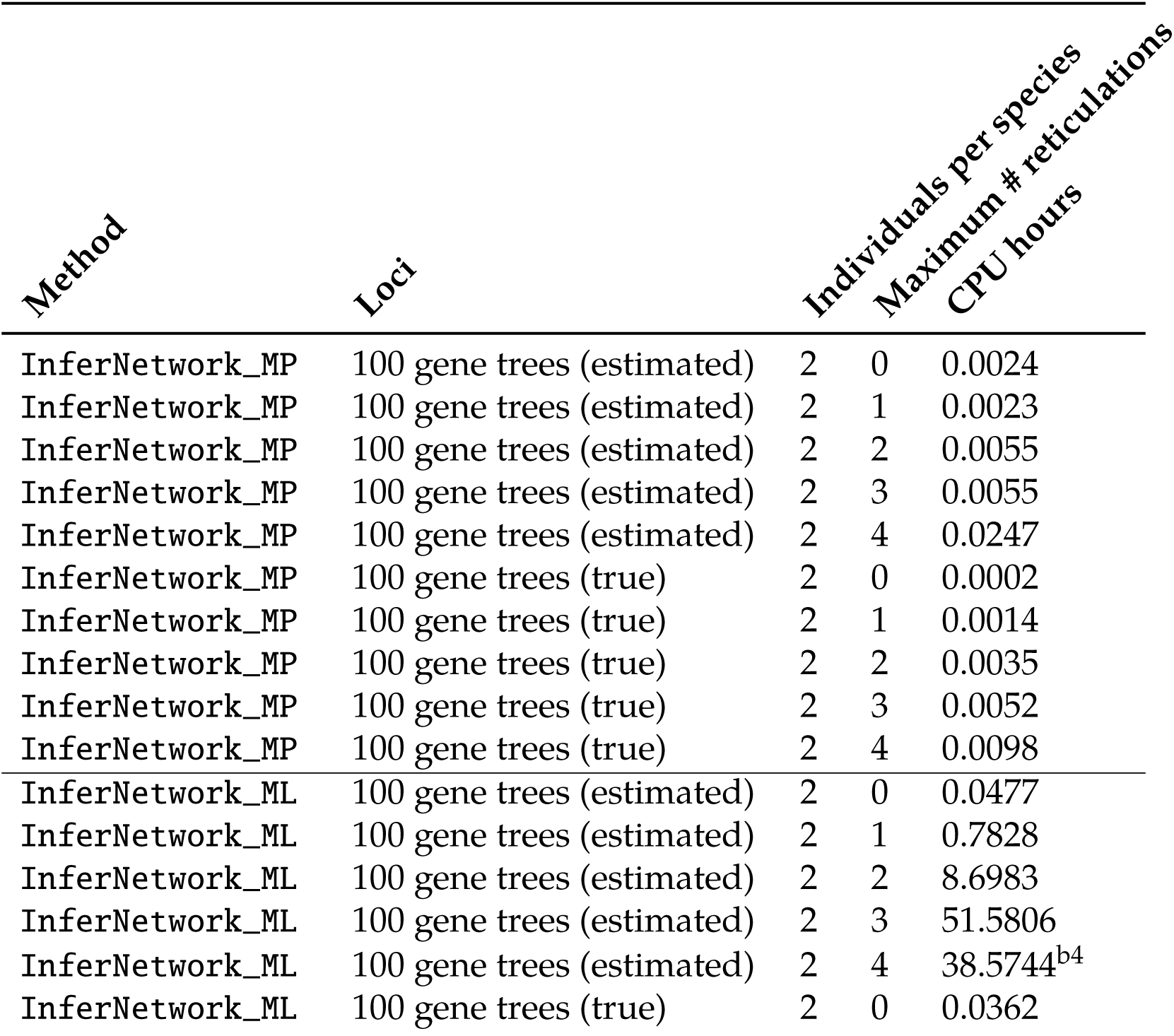
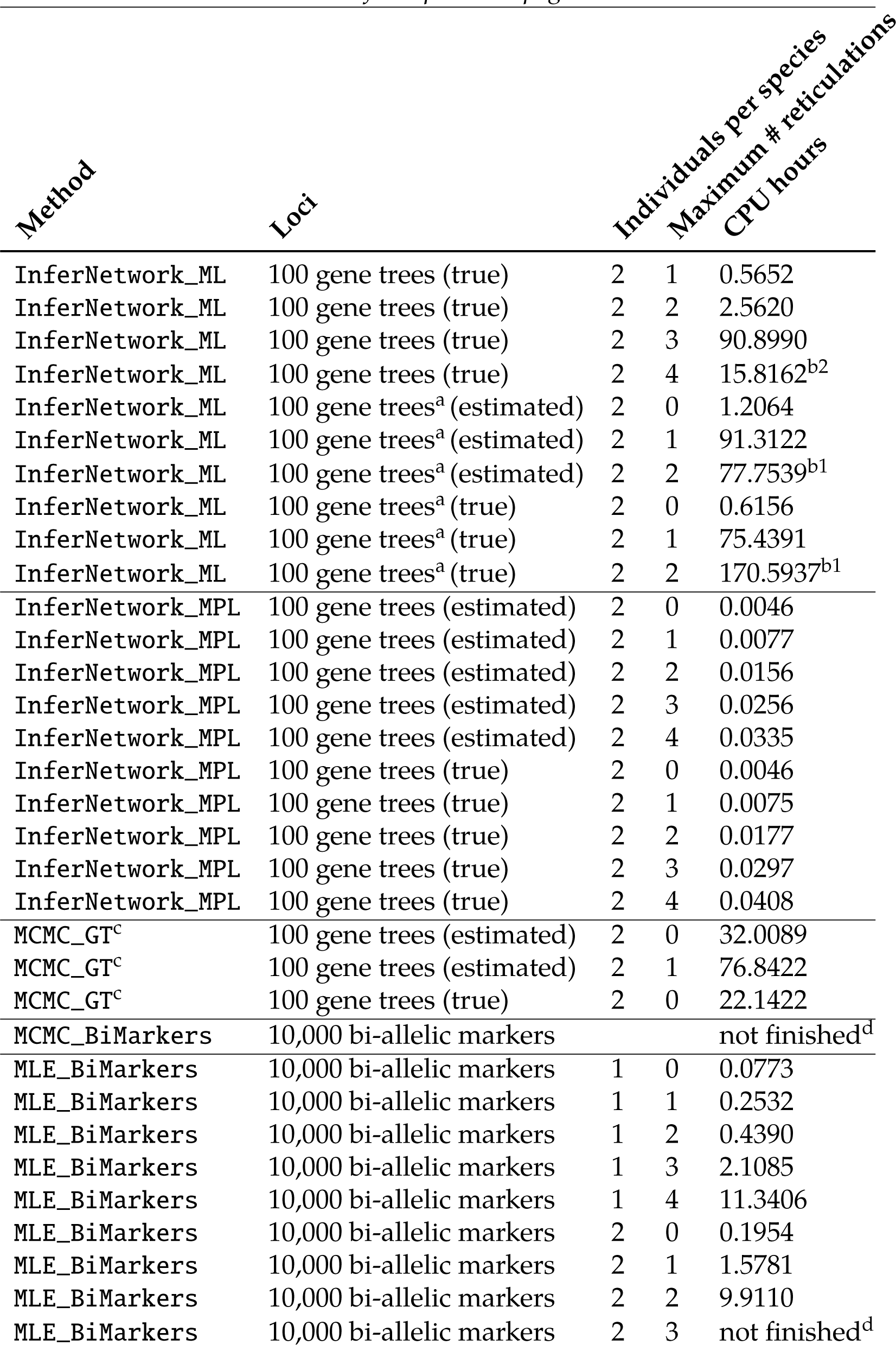
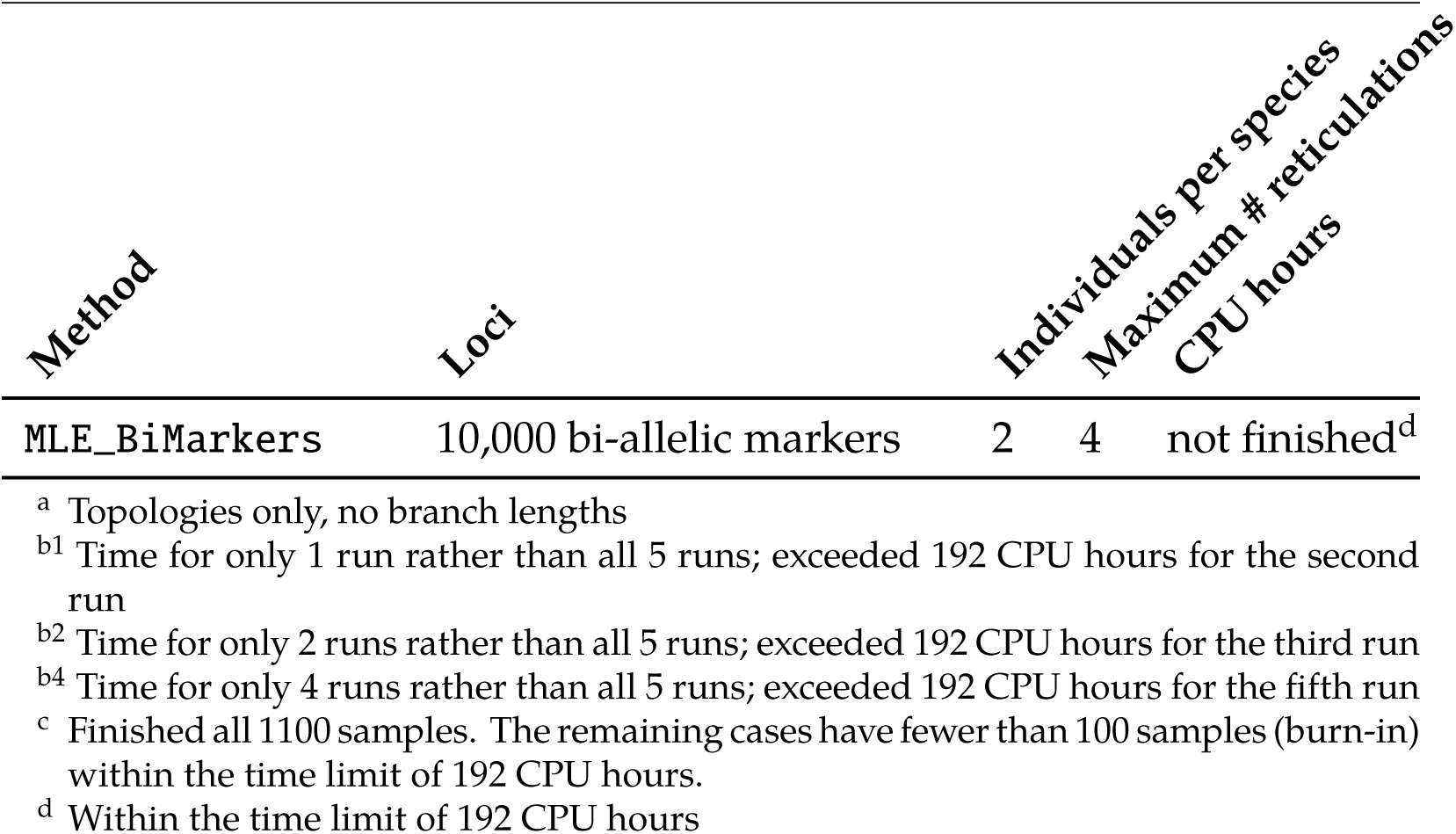
Computational requirements for methods other than `MCMC_SEQ` on the data set and analyses reported above in this section.

## 5 Analyzing Larger Data Sets

Statistical inference of phylogenetic networks is computationally very demanding, as the computational requirements of likelihood calculations on networks can be orders of magnitude larger than those on trees [47, 8]. Currently, analyses of larger data sets in terms of the numbers of loci and taxa can be done in PhyloNet using one of three ways:

1. **Maximum pseudo-likelihood inference:** Inference using maximum pseudo-likelihood from gene tree topologies (the `InferNetwork_MPL` command). The advantage of this approach is that computing the pseudo-likelihood of a network candidate is very efficient (can be done in seconds or minutes even on networks with 200 taxa). However, this approach does not circumvent the problem of searching the enormous space of phylogenetic networks. Therefore, maximum pseudo-likelihood inference of very large networks can suffer in practice from getting stuck in local optima. Furthermore, while pseudo-likelihood performed well on the data set above in terms of determining the number of reticulations, this is not the case in general.
2. **Tree-based augmentation:** Inference of a species tree and then augmenting it into a phylogenetic network (the `InferNetwork_MPL -fs` or `InferNetwork_MDC -fs` command). The advantage of this approach is that there are very efficient methods for inference of species trees, and then the number of ways to add *k* reticulations to the tree is polynomial in *n*. The disadvantages of this approach are that the start tree could be wrong, thus resulting in a wrong network. Furthermore, most criteria for evaluating the network candidates are computationally very demanding, leaving mainly the MDC and pseudo-likelihood criteria to use. In other words, this can be viewed as a very constrained version of 1 where the search space consists of only augmentation of a given tree into a network. The merits and limitations of this approach were recently studied in [1].
3. **Divide-and-conquer:** Inference based on a divide-and-conquer approach (the `NetMerger` command). This approach is based on the recently developed divide-and-conquer method of [46], where the set of taxa is divided into overlapping 3-taxon subsets, trinets (networks on three taxa) are inferred, and then these trinets are merged into the full network. A major advantage of this method is that it avoids searching the space of large phylogenetic networks, as only the space of trinets is searched (in parallel for each 3-taxon subset). Furthermore, model complexity is handled in the inference of the trinets; combining them together need not deal with it.

The networks in Fig. 1 and Fig. 2 are in fact sampled from the larger network shown in Fig. 15(a) whose inference is infeasible via direct application of `InferNetwork_MP`, `InferNetwork_ML`, `MCMC_GT`, `MCMC_SEQ`, `MCMC_BiMarkers`, or `MLE_BiMarkers`.^1^

**Figure 15:**
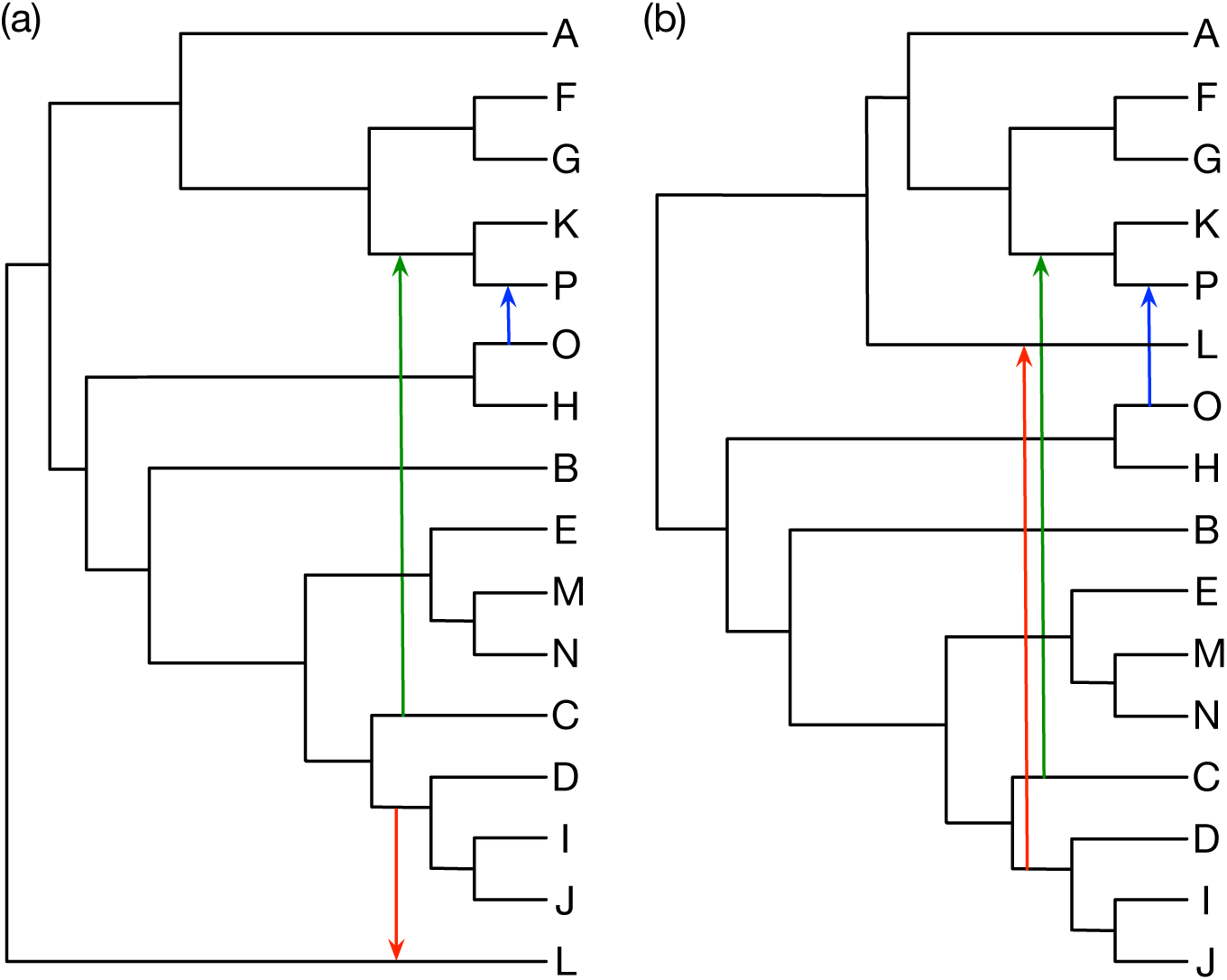
Analysis of a larger phylogenetic network. (a) A model phylogenetic network on 16 taxa with outgroup taxon *Z*. (b) The phylogenetic network inferred by the divide-and-conquer method of [46].

Maximum pseudo-likelihood inference using the `InferNetwork_MPL` command using the gene trees estimated by IQTREE and setting the maximum number of reticulations at 3 yields the correct network that is shown in Fig. 15(a). The log pseudo-likelihood of the network is -158498.

For the tree-based augmentation approach, we ran ASTRAL [45] on the gene trees (estimated by IQTREE) to infer a backbone tree. ASTRAL inferred the correct backbone tree; that is, the tree obtained from the network in Fig. 15(a) by removing the three arrows. The method then searched for augmentation of the tree into a network in four different ways: searching for the optimal network under the pseudo-likelihood criterion of [43] and under the MDC criterion of [40], when the number of reticulations was set at 3 and at 5 for each of the two criteria. Augmentation into networks based on both criteria resulted in the correct network when the number of reticulations was set at 3. The log pseudo-likelihood and MDC score of the resulting network are -158498 and 173, respectively. However, when the number of reticulations was set at 5, both criteria resulted in two additional reticulations (the true network with three reticulations, plus two additional reticulations, for a total of five reticulations) with log pseudo-likelihood and MDC score of -158483 and 160, respectively. Once again, these results illustrate that inference based on these criteria yields good results, but that both of them suffer from the problem of determining the correct number of reticulations. Finally, using the divide-and-conquer method of [46], the inferred network is shown in Fig. 15(b). As the result shows, the only error in the inferred network is the placement of taxon L. However, despite this wrong placement, the reticulation involving L and the clade (D,(I,J)) is inferred correctly. It is important to highlight here that the number of reticulations need not be specified as the trinets were inferred using the Bayesian method `MCMC_SEQ` which determined the number of reticulations in each trinet individually. Furthermore, it is worth mentioning that when the true trinets were used in `NetMerger`, the correct network was inferred, which points to inaccuracy in some of the inferred trinets as the cause of the wrong placement of L.

With respect to the running time, maximum pseudo-likelihood inference took 13.8293 CPU hours for 10 runs when the maximum number of reticulations was set to 3. Under the pseudo-likelihood criterion, the tree-based augmentation approach took 7.2995 and 10.3108 CPU hours when the maximum number of reticulations was set to 3 and 5, respectively, for 10 runs. Under the MDC criterion, when the number of reticulations was set to 3 and 5, the tree-based inference took 3.9351 and 24.4727 CPU hours, respectively, for 10 runs. The divide-and-conquer inference took 1337.8253 CPU hours inferring the 680 3-taxon subnetworks by `MCMC_SEQ` (this can be trivially parallelized, but the time we report is the total time for inferring all 680 3-taxon subnetworks sequentially), while it took only 2.34 minutes for the merger procedure of obtaining the full network from the 680 3-taxon subnetworks.

## 6 Comparing and Summarizing Networks

While comparing and summarizing trees is easy, the same tasks are very hard for networks. For example, while testing if two trees are isomorphic can be done in polynomial time, the problem is NP-hard for phylogenetic networks in general. This is why various dissimilarity measures for restricted classes of phylogenetic networks have been devised [22, 3, 4, 2, 11, 5, 36]. While PhyloNet has utilities to compare phylogenetic networks based on their constituent trees, tripartitions, and clusters, it also has a function that computes the distance measure of [22]. However, all these measures are very sensitive and return “inflated” dissimilarity values even for what would, visually, be considered a small difference between the networks. We now discuss five features in PhyloNet for summarizing sets of phylogenetic networks with the goal of elucidating common substructures in sets of networks that are obtained by an inference method. The input to a summarization method is a set of networks obtained by an inference method (e.g., a set of equally optimal networks under the MDC criterion, or a set of the different network topologies collected from MCMC). The output is a summary that differs depending on the method used, as we now describe. The input to the summary task is a set of networks {Ψ_1_, Ψ_2_, …, Ψ_*n*_}, and the output is a set of pairs {(*S*_1_, *v*_1_), (*S*_2_, *v*_2_), …, (*S*_*m*_, *v*_*m*_)}, where each *S*_*i*_ is a substructure obtained from the networks and *v*_*i*_ is its frequency in the input networks (normalized by the number of networks so that all frequencies add up to 1).

### 6.1 Displayed Trees

A network with *k* reticulations displays up to 2^*k*^ trees, where each tree is obtained from the network by removing one of the two reticulation edges for each reticulation node. The summary based on displayed trees consists of the union of the displayed trees of all networks and the frequency of each tree is derived based on the number of networks by which it is displayed.

### 6.2 Backbone Networks

Backbone networks extend the notion of displayed trees. Given a network Ψ with *k* reticulations, a backbone of it is a network Ψ′ that is obtained by removing one of the two reticulation edges for each node in a *subset* of the set of all reticulation nodes. If the subset contains all reticulation nodes, the resulting backbone is also a displayed tree. If the subset is the empty set, the resulting backbone network is the network Ψ itself. Thus, there are up to 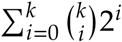 backbone networks. The summary based on backbone networks consists of the union of the backbones of all networks and the frequency of each backbone is derived based on the number of networks of which it is a backbone.

### 6.3 Tree Decompositions

As network can be decomposed into a forest of trees by applying the following operation to each reticulation node *u* whose parents are *v*_1_ and *v*_2_:

- Create two nodes *x*_1_ and *x*_2_ and label both of them by a unique name, say, Q;
- Delete the two edges (*v*_1_, *u*) and (*v*_2_, *u*); and,
- Add two edges (*v*_1_, *x*_1_) and (*v*_2_, *x*_2_).

This operation was illustrated in Fig. 3 in [33]. Given the tree decomposition of all networks in the input, a summary consists of all the unique trees in the union of resulting forests, along with the frequency of each tree.

### 6.4 Tripartitions

Each reticulation node *u* can be represented in the form of tripartition as *X*: *P*_1_ | *P*_2_, where *X* is the set of taxa that are under this reticulation node, and *P*_1_ and *P*_2_ are two sets of taxa that are under the two siblings of this reticulation node, respectively. When one of the parents of the reticulation node *u* is also a reticulation node, *u* has only one sibling, and the tripartition is of the form *X*: *P*. When both parents of *u* are reticulation nodes, *u* has no siblings, and the tripartition is of the form *X*. For example, for the reticulation node denoted by the green circle in Fig. 1, the tripartition is

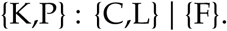

The rationale behind a tripartition is that it illustrates that the clade containing the leaves in *X* is a hybrid, and that the two clades given by the sets of taxa *P*_1_ and *P*_2_ are the parents of the hybrid. Once the tripartitions of all networks in the input are computed, they are displayed along with their frequencies across all networks (each unique tripartition appears once).

### 6.5 Major Trees

For inference methods that produce networks with estimated inheritance probabilities, the major tree of a network is the one that is obtained from the network by removing for each reticulation node the reticulation edge with the lower inheritance probability (one is chosen arbitrarily if the inheritance probability is 0.5). The summary, in this case, consists of all major trees of the networks (each unique tree appears once) and for each the number of networks of which it is the major tree.

## 7 What Reticulate Evolutionary Processes Does PhyloNet Handle?

We believe the proliferation in the literature of specialized terms of *reticulate evolution* hampers, rather than helps, the development of general tools for inferring reticulate evolutionary histories. It is unclear, for example, what the difference between hybrid speciation and hybridization is from a modeling point of view. While the former results in a new species and the latter does not, from a modeling and inference perspective that distinction seems irrelevant. Gene flow has been defined as the “exchange of genes between two populations as a result of interbreeding” whereas admixture is “a sudden increase in gene flow between two differentiated populations” [29]. But, it is unclear if these distinction and naming have any implications on developing computational methods for inference. As far as PhyloNet application is concerned, all these processes are collectively referred to as reticulation; elucidation of the actual biological process that gave rise to the reticulation is not within the capabilities of phylogenetic methods.

Let us illustrate the issue with the phylogenetic networks in Fig. 16. What is shown is *one* phylogenetic network that is visualized in two different ways. That is, in graph-theoretic jargon, the two networks shown in Fig. 16 are *isomorphic*. Presented with multi-locus data from the three taxa A, B, and C, the various inference utilities in PhyloNet yield a single phylogenetic network (let us assume here that the inferred network is unique). Calling the reticulation event hybridization and drawing the network as in Fig. 16(a) or calling the event hybrid speciation and drawing the network as in Fig. 16(b) is not a distinction made by PhyloNet nor is it an aspect that is inferable from data. Such a distinction can be made after the network is inferred, and using some knowledge that is “external” to the inferred network topology.

**Figure 16:**
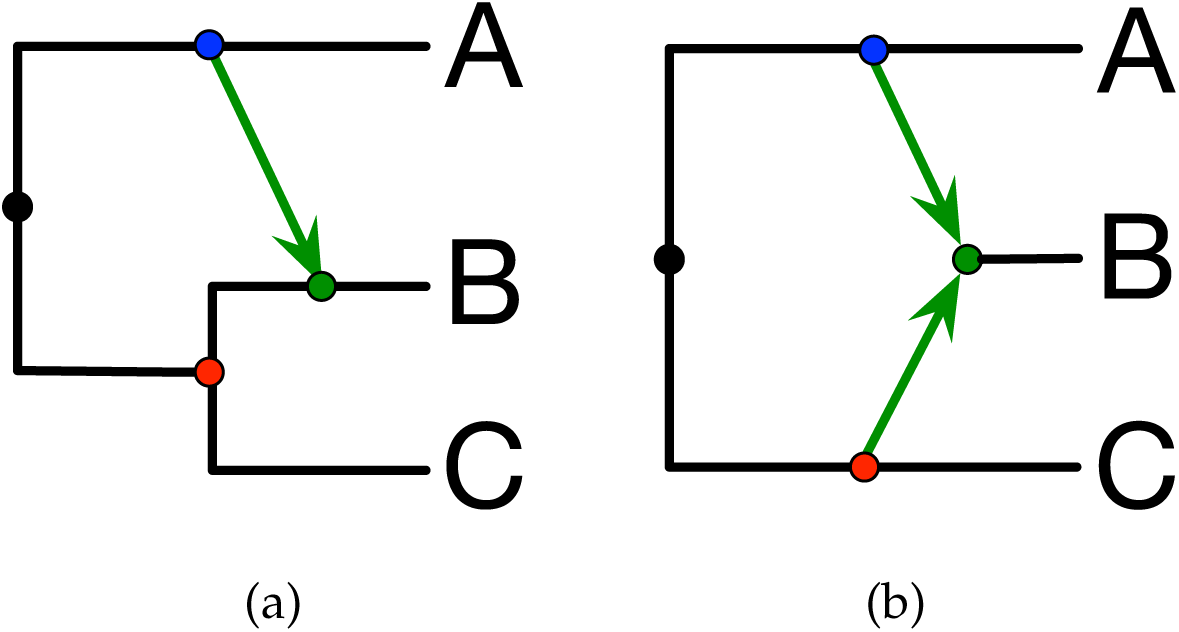
Visualizing the same network in different ways conveys different biological stories. (a) The phylogenetic network is drawn to convey the message that (A,(B,C)) is a species tree, and that there is hybridization involving an ancestor of A and an ancestor of B. (b) The phylogenetic network is drawn to convey that hybrid speciation occurred between (ancestors of) species A and C and gave rise to new species B. Nodes with the same color in the two networks correspond to each other.

Along the same lines, if A, B, and C were different species, the reticulation event illustrated in Fig. 16 would be referred to as hybridization or hybrid speciation. On the other hand, if the same exact network was inferred from data collected from different individuals collected from sub-populations of the same species, the reticulation event would be referred to as gene flow or admixture. And to hammer the point home even further, if the data consists of a sequence alignment from three individuals in one population and recombination was present, the phylogenetic network of Fig. 16 would be referred to as an *ancestral recombination graph* (ARG) and the reticulation node would denote a recombination event.

Therefore, it is important to distinguish between the networks that are inferred by methods such as those in PhyloNet, where the non-treelike events should be generically referred to as reticulation, and the actual biological process that took place (e.g., hybridization vs. hybrid speciation vs. admixture) and whose elucidation is not within the realm of what a phylogenetic inference method is capable of. This very important point is a major theme in the excellent book on phylogenetic networks by David Morrison [20]. As Morrison wrote, “Unfortunately, there is little that a mathematical network can necessarily tell us about any underlying biology. That a particular taxon is apparently the descendant of a reticulation event does not actually tell us what that event was, nor even whether the correct taxon has been identified. For example, how to distinguish HGT and hybridization mathematically is not obvious.” A method’s inability to distinguish, say, horizontal gene transfer from hybridization, is not a weakness or limitation of the method. To the contrary, a claim that a method can make such a distinction from molecular data should raise questions.

Why, then, emphasize the process of hybridization with respect to PhyloNet, including in the titles of manuscripts as in [41, 40, 8]? The reason has to do with the model employed by PhyloNet and the detectability of reticulation. While a thorough study is currently lacking, we believe the signal in data obtained from a single individual within each sub-population is not sufficient to infer the phylogenetic network that represents the network-like demographic structure of the entire population. This is why the more popular methods in population genetics, namely STRUCTURE [30] and TreeMix [28], rely on allele frequency data that is obtained from many individuals within the population. Similarly, we believe all the complexities that arise in prokaryotic evolution due to processes such as gene fusion and fission [19], coupled with the fact that in some cases only a single gene could be transferred make it very challenging and potentially not useful to apply current functionalities in PhyloNet to analyze the evolutionary history of prokaryotic genomes.

### 7.1 How Can I Analyze Polyploids?

Despite the fact that polyploidization plays an important role in plant evolution, few studies attempt to specifically handle reticulations arisen from polyploidy, especially in the presence of ILS. Lott *et al.* [15] presented a method to construct a consensus tree by summarizing a collection of multi-labeled gene trees. Jones *et al.* [13] developed a statistical framework that could accurately infer the evolutionary history of allopolyploids but restricted to networks of diploids and tetraploids. Gregg *et al.* [9] developed an algorithm to detect the polyploidy events by reconciling the known (multi-labeled) species tree and a set of (multi-labeled) gene trees but not accounting for ILS. Recently, Oberprieler *et al.* designed a workflow for reconstructing the reticulate evolutionary history involving polyploid complexes from multi-locus data using MDC-based inference in PhyloNet. The workflow uses a permutation approach to assign homoeologs in polyploids and then constructs the multi-labeled species tree under the MDC criterion. The algorithm proceeds in the following steps:

1. For each polyploid species and each gene tree, it generates all possible diploid parental inheritance histories of all possible combinations of allele pairs. To determine the polyploid-specific and locus-specific optimal allele pairs inheritance history, the MDC score is computed by running the `Infer_ST_MDC` command for each of the possible histories, and the one with minimum MDC score is kept.
2. Based on the polyploid-specific and locus-specific optimal allele combinations at all sequenced loci generated from step 1, it uses the `Infer_ST_MDC` command to compute the MDC score and find the polyploid-specific optimal allele combinations across loci.
3. After obtaining the polyploid-specific optimal allele combinations across loci for all polyploids, it runs the `Infer_ST_MDC` command to reconstruct the overall MUL-species tree and finally converts the resulted MUL-species tree into species network.

However, such a permutation approach can be executed directly using the `InferNetwork_MP` command. The `InferNetwork_MP` command would compute the optimal MDC score by converting the species network into the corresponding MUL-tree. Then all possible allele mappings to the leaves of the MUL-tree are produced and the mapping that gives rise to the minimum MDC score is selected as the optimal coalescent history. Through this way, the optimal assignment of homoeologs to parental lineages in polyploids based on the MDC criterion is obtained. To illustrate this approach, we reanalyzed the *Leucanthemopsis* test data set of [24] which consists of 12 diploid, tetraploid and hexaploid representatives of *Leucanthemopsis* to reconstruct reticulate evolution involving hybridization and polyploidization, and compared the result from the `InferNetwork_MP` command to that from the permutation approach through the `Infer_ST_MDC` command [24]. It is important to note that the given data set only consists of 5 inferred gene trees and that this small number of loci might not be sufficient to reconstruct a reliable history.

To construct the reticulate evolutionary history, we set the maximum number of allowed reticulations to 0, 1, 2, and 3. With increasing number of reticulations from 0 to 3, the sequence of MDC scores obtained was 139, 136, 135, and 135, respectively; the networks are shown in Fig. 17. It took less than 3.5 minutes to infer each of the resulting networks over 20 runs using one processor. Observe that the improvement in the MDC score from the 0 reticulations to the one with one/two reticulation(s) is very small. And when setting the maximum number of allowed reticulations to 3, two of the three returned species networks with equivalent optimal MDC score only have 2 reticulations. This observation shows that it is very unlikely to have more than one reticulation for this data set (given the five gene trees used). One major difference between the species network with 0 reticulations and with 1 reticulation is that the tetrapolyploid *L. alpina* is found to have autopolyploid origin in the former whereas it is assumed to be allopolyploid and formed from the hybridization between *L. pallida* subs. *virescent* var. *bilbilitanum*. The inferred species network from the permutation approach yields an MDC score of 155 (the `DeepCoalCount_network` command), which is greater than the one with 0/1/2/3 reticulations found by `InferNetwork_MP`.

**Figure 17:**
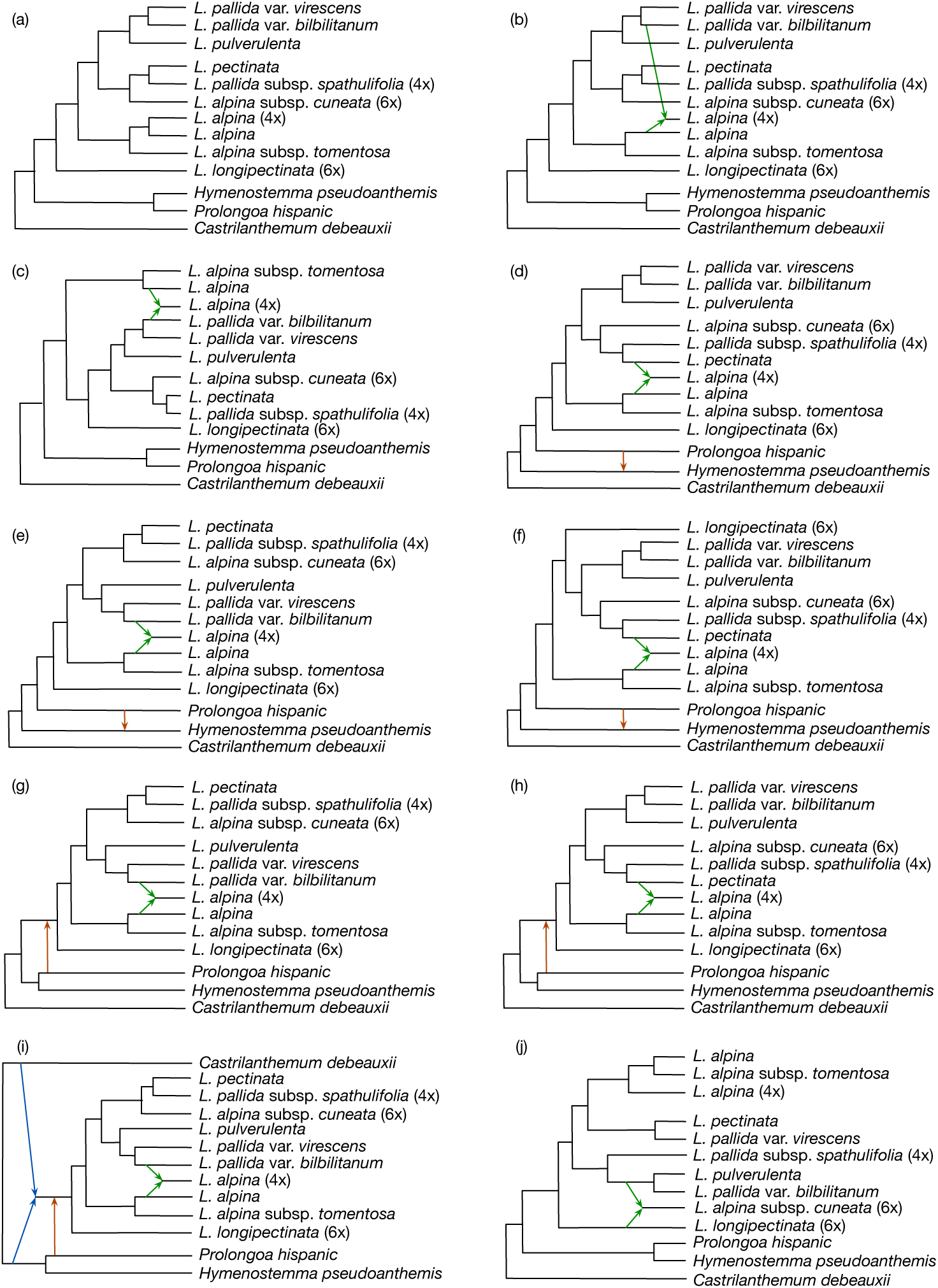
Inference results using the MDC criterion on the example data set of *Leucanthe-mopsis* representatives in [24]. (a)-(i) Species networks returned by `InferNetwork_MP`. (a), (b)-(c), (d)-(f) and (g)-(i) are optimal networks inferred when the maximum number of reticulations was set to 0, 1, 2 and 3, respectively, and the corresponding optimal MDC scores are 139, 136, 135, and 135, respectively. (j) The phylogenetic network inferred by the permutation approach of [24] and whose MDC score is 155 as computed by the `DeepCoalCount_network` command.

If some species are known to be hybrid, `InferNetwork_MP` allows the user to restrict the search space of the species networks by forcing a set of species to be hybrid species using the `-h` option. We ran the analysis on the same data set, with one specified hybrid species *L. alpina* subsp. *cuneata*, and with two specified hybrid species, namely *L. alpina* subsp. *cuneata* and *L. longipectinata*, separately. The inferred species networks are shown in Fig. 18. For the former case, we ran the `InferNetwork_MP` with the maximum number of allowed reticulations set to 1, 2 and 3, while for the latter one, the maximum number of allowed reticulations was set to 2 and 3. When only *L. alpina* subsp. *cuneata* was forced to be a hybrid species, the inferred species networks are of equivalent MDC score 143 (Fig. 18(a)(b)) and only one reticulation event was identified no matter what the maximum number of allowed reticulations was. When it comes to two specified hybrid species, with maximum number of allowed reticulations set to 2 and 3, unique species networks, Fig. 18(c) with MDC score 138 and Fig. 18(d) with MDC score 136, respectively, were returned. Why searching in a more restricted phylogenetic network space would result in a network topology with better MDC score? It is worth repeating the message above that `InferNetwork_MP` uses the hill climbing strategy to search for the optimal network. When only one hybrid species *L. alpina* subsp. *cuneata* was specified, in the limited number of runs, `InferNetwork_MP` had not searched over the more restricted space (i.e., the search space of two specified hybrid species *L. alpina* subsp. *cuneata* and *L. longipectinata*) yet, and thus it did not find a network with the same or better MDC score.

**Figure 18:**
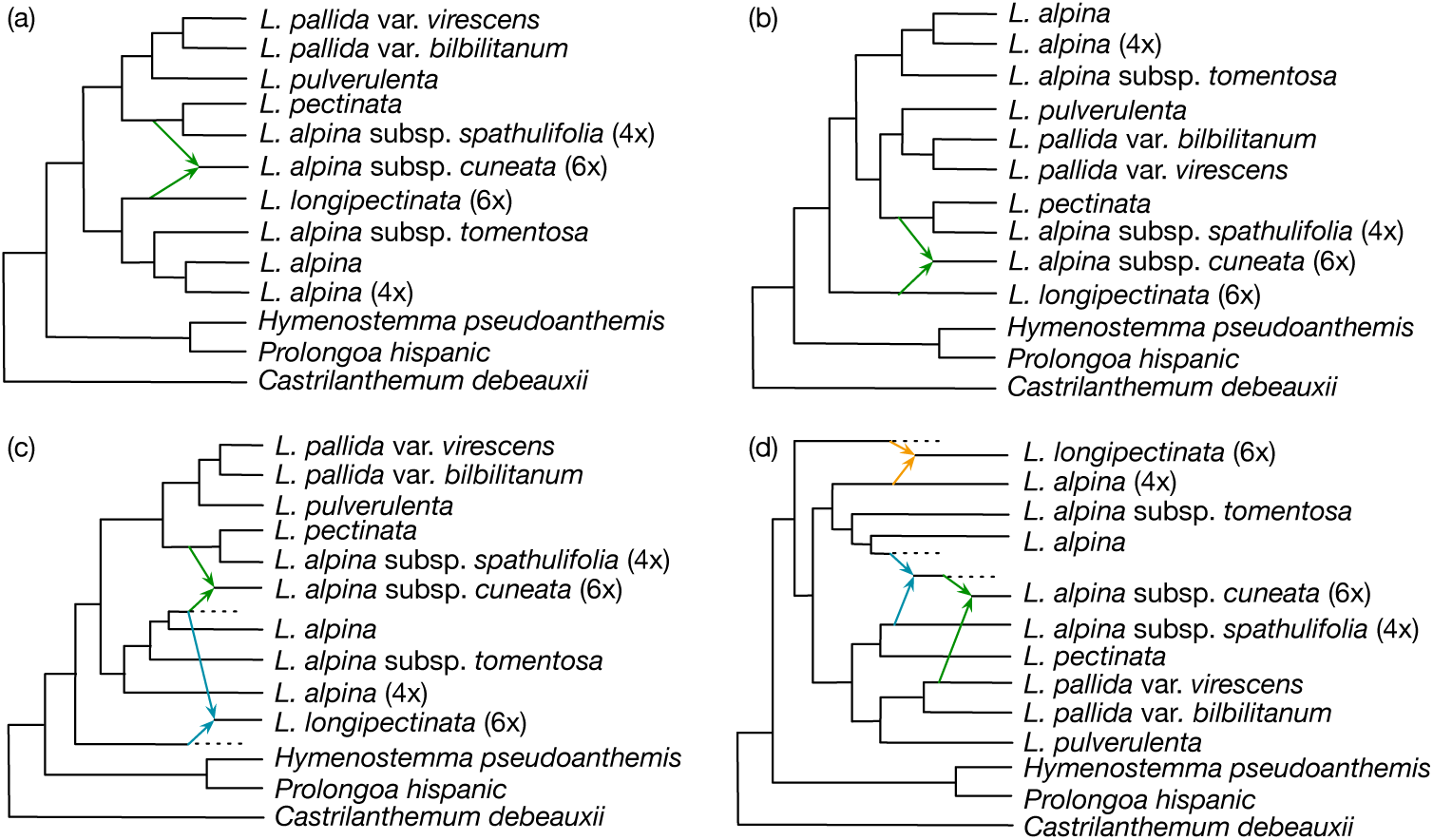
Inference results using the MDC criterion on the example data set of *Leucanthe-mopsis* representatives in [24] with specified hybrid species. When forcing the hexapolyploid *L. alpina* subsp. *cuneata* to be a hybrid species with the `-h` option, and setting the number of reticulations to 1, 2 and 3, `InferNetwork_MP` returned two distinct species networks (a,b) with identical MDC scores of 143. When specifying the hexapolyploid *L. alpina* subsp. *cuneata* together with the hexapolyploid *L. longipectinata* to be hybrid species, and setting the number of reticulations to 2 and 3, `InferNetwork_MP` inferred a network with a MDC score of 138 (c), and a network with a MDC score of 136 (d), respectively.

MUL-trees were used in [41, 40] as the representation on which likelihood and parsimony computation were done. A thorough analysis of the connection between MUL-trees and phylogenetic networks, including cases of uniqueness (a MUL-tree uniquely defines a phylogenetic network) is given in [10]. As such, in addition to `InferNetwork_MP`, other methods in PhyloNet could be used to analyze data sets with polyploids.

Finally, while we demonstrated how methods in PhyloNet can be used to analyze data with polyploids, much work is still needed to accurately model the complexity that arises in the presence of polyploids, in particular when the mode of polyploidy—allo or auto—needs to be elucidated as one is about reticulation and the other is effectively about whole-genome duplication. The recent review of Oxelman *et al.* [26] discusses existing methods and their applications to data with polyploid species, as well as challenges and future directions in the field.

## 8 Conclusions

Our group has been developing the PhyloNet software package for over 10 years. Our ultimate goal is to contribute to the development of a wide array of tools that will make phylogenetic network analysis and use in evolutionary biology as simple and natural as that of phylogenetic trees.

In this paper, we discussed practical aspects that relate to data analyses using PhyloNet. This software package has been developed to enable evolutionary analyses of data sets where reticulation is suspected, and consists of several inference functionalities that employ different criteria and inference algorithms. Analyzing and interpreting phylogenetic networks is much more challenging than analyzing and interpreting trees; therefore, it is important that the results obtained by network inference methods are inspected carefully. A major challenge in this area is determining the true number of reticulations that had occurred during the evolutionary history of the taxa under consideration. While the Bayesian approach allows for incorporating model complexity in a natural way via the prior distribution, extra caution must be taken when inferring networks using any of the inference methods in order to distinguish between real reticulations and false ones whose addition is simply a case of making von Neumann’s elephant wiggle its trunk.^1^

While PhyloNet is capable of inferring phylogenies with reticulations, it cannot determine the biological processes behind those processes. One major direction for research and addition of features to PhyloNet relates to modeling gene duplication and loss—in addition to reticulation and ILS, which are currently handled by the software package. Furthermore, PhyloNet neither makes assumption about species nor delimits them. At an abstract level, whatever taxa the user specifies as different species, PhyloNet handles them as such.

The relationship between species networks and trees is intricate. While networks extend trees, inferring a network with no reticulations—that is, a tree—does not necessarily mean a correct tree (whatever that tree means) is inferred. The flip side of this coin is that network inference by first inferring a tree and then “completing” it into a network is not necessarily a good approach [1]. Methods implemented in PhyloNet infer evolutionary histories by directly searching the space of phylogenetic networks, and not the space of completions of a given tree into a network, though we provide such a feature for those interested in using it.

Last but not least, computational requirements of phylogenetic network inference remain the major bottleneck for almost all methods in PhyloNet. Direct applications of maximum parsimony, maximum likelihood, and Bayesian inference to large data sets, especially in terms of the number of taxa, could be infeasible. Inferences based on pseudo-likelihood functions and divide-and-conquer approaches are very promising in such cases and are enabled by PhyloNet.

## Acknowledgments

Beside the authors on this paper, several former members of the group of L.N. contributed significantly to the development of PhyloNet, listed in chronological order of their time in the group: Derek Ruths, Cuong V. Than, Hyung Jung Park, Yun Yu, Nikola Ristic, Jianrong Dong, Kevin Liu, Dan Ye, Dingqiao Wen, Peng Du, Ryan (Leo) Elworth, Jiafan Zhu, and Hunter Tidwell, and Yaxuan Wang. Furthermore, the development of the likelihood model in PhyloNet was done in collaboration with Prof. James H. Degnan.

The development of PhyloNet has been ongoing for over 10 years and has been generously supported by several grants and awards from various agencies and foundations. We acknowledge all support—prior and current—that has facilitated this development: DOE grant DE-FG02-06ER25734, NIH NLM grant R01LM009494, NSF grants CCF-0622037, DBI-1062463, CCF-1302179, DBI-1355998, CCF-1514177, CCF-1800723, and DMS-1547433, and fellowships to L.N. from the Alfred P. Sloan Foundation and the John Simon Guggenheim Memorial Foundation.

## Footnotes

1 Mathematically, the definition of an inheritance probability is not this simple, as each locus could have its own inheritance probability, as defined and discussed in detail in [42]. Defining one inheritance probability for each locus and for each reticulation edge poses significant challenges in terms of inference, which is why all functionalities in PhyloNet assume a single inheritance probability across all loci per reticulation node. An alternative is to integrate out these probabilities during inference, which can be done analytically [35].

2 One could use methods in the second category in an optimization setting by returning from all samples collected the one with the highest posterior, which would be the *maximum a posteriorio*, or MAP, estimate.

3 For example, a drop in the MDC score from 4 to 3 is a 25% improvement, but that is just an artifact of the small denominator value of 4.

4 While `MLE_BiMarkers` utilizes pseudo-likelihood computations, these can be computationally very expensive on large networks since full likelihood calculations are carried out on subnetworks.

5 According to Enrico Fermi, the famed mathematician John von Neumann said “With four parameters you can fit an elephant to a curve; with five you can make him wiggle his trunk.”

